# Rational Tuning of CAR Tonic Signaling Yields Superior T-Cell Therapy for Cancer

**DOI:** 10.1101/2020.10.01.322990

**Authors:** Ximin Chen, Mobina Khericha, Aliya Lakhani, Xiangzhi Meng, Emma Salvestrini, Laurence C. Chen, Amanda Shafer, Anya Alag, Yunfeng Ding, Demetri Nicolaou, Junyoung O. Park, Yvonne Y. Chen

## Abstract

Chimeric antigen receptors (CARs) are modular proteins capable of redirecting immune cells toward a wide variety of disease-associated antigens. Here, we explore the effects of CAR protein sequence and structure on CAR-T cell function. Based on the empirical observation that CD20 CARs with similar sequences exhibit divergent tonic-signaling and anti-tumor activities, we devised engineering strategies that aimed to improve CAR-T cell function by tuning the intensity of tonic signaling. We found that CARs designed to exhibit low but non-zero levels of tonic signaling show robust effector function upon antigen stimulation while avoiding premature functional exhaustion by CAR-T cells. Through alterations of the CAR’s ligand-binding domain and overall protein conformation, we generated CD20 CAR variants that outperform the CD19 CAR in mouse models of human lymphoma. We further demonstrate that rational modification of protein confirmation can be generalized to improve GD2 CAR-T cell efficacy against neuroblastoma. These findings point to tonic signaling and basal T-cell activation as informative parameters to guide the rational design of next-generation CARs for cancer therapy.

## INTRODUCTION

The adoptive transfer of chimeric antigen receptor (CAR)-T cells has shown remarkable efficacy for advanced B-cell malignancies, and CAR-T cell candidates against a wide variety of cancer types have entered clinical testing. However, few have shown comparable anti-tumor efficacy to CD19 CAR-T cells to date (Grigor et al., 2019; Guedan et al., 2018; Majzner and Mackall, 2019; Yamamoto et al., 2019). Standard approaches to CAR construction rely on recombining a limited set of extracellular spacer, transmembrane, and cytoplasmic signaling domains with ligand-binding moieties specific to the antigen of interest. While this empirically driven process can reliably yield CAR molecules with tumor reactivity, the resulting CAR-T cells often fail to achieve robust clinical efficacy. For example, although CD20 CAR-T cells showed promising activity in preclinical models, clinical trial results have not yet achieved similar efficacy compared to CD19 CAR-T cell therapy against the same disease types (Till et al., 2012; Zhang et al., 2016). Similarly, a second-generation GD2 CAR incorporating the CD28 co-stimulatory domain has been shown to exhibit inferior *in vivo* anti-tumor function compared to a CD19 CAR containing the same signaling domains (Long et al., 2015). The inability to consistently generate robustly functional CARs against antigens of interest highlights the need to further understand the molecular and mechanistic relationship between CAR design and CAR-T cell function.

CAR engineering efforts thus far have revealed several design parameters that influence CAR-T cell function (Chang and Chen, 2017; Hong et al., 2020). These include co-stimulatory domains that booster T-cell activation upon antigen stimulation (Omer et al., 2018; Wijewarnasuriya et al., 2020; Zhao et al., 2015), extracellular domains that provide structural support for optimal T-cell/target-cell conjugation (Hudecek et al., 2013; Srivastava and Riddell, 2015), binding affinity between the CAR and the targeted antigen (Drent et al., 2019; Liu et al., 2015), as well as CAR-independent parameters such as antigen expression level on target cells (Majzner et al., 2020; Watanabe et al., 2015). However, these parameters cannot fully explain the difference between the CD19 CAR and other constructs such as CD20 CARs, which have similarly high binding affinities, contain the same co-stimulatory domains, and bind an antigen that is also highly expressed on the same type of tumor cells as targeted by the CD19 CAR.

More recently, several studies highlighted the phenomenon of CAR tonic signaling—i.e., CAR signaling in the absence of antigen stimulation—and its potential role in triggering premature T-cell dysfunction (Frigault et al., 2015; Gomes-Silva et al., 2017; Long et al., 2015; Watanabe et al., 2016). It has been suggested that the robust functionality of CD19 CAR-T cells can be attributed to a lack of tonic signaling by the CD19 CAR (Frigault et al., 2015; Long et al., 2015). However, phenotypic descriptions of tonically signaling T cells have been varied, and hypotheses on what specific CAR components could cause or prevent tonic signaling are at times contradictory (Frigault et al., 2015; Gomes-Silva et al., 2017; Long et al., 2015).

Here, we report that CAR sequence and architecture have significant and tunable impacts on tonic signaling, and that CAR-T cell metabolism and anti-tumor function can be altered through the tuning of tonic-signaling intensity and associated basal T-cell activation. We show that a low but non-zero level of tonic signaling can increase CAR-T cell function by potentiating rapid anti-tumor response while avoiding premature T-cell exhaustion, and the level of basal T-cell activation can be tuned through the insertion of torsional linkers and the sequence hybridization of different ligand-binding domains. By applying these protein-engineering strategies, we generated novel CD20 CARs that outperform the CD19 CAR in mouse models of B-cell lymphoma, and found memory phenotype enrichment and minimization of CAR-driven metabolic disturbance as properties associated with improved CAR-T cell function. These results point to the rational tuning of tonic signaling and basal T-cell activation as a useful and potentially broadly generalizable approach to engineering robust CAR-T cell therapies for cancer.

## RESULTS

### scFv sequence alters tonic signaling and CAR-T cell metabolism

To date, studies on tonic signaling have primarily relied on comparing CARs targeting different antigens, thus complicating the interpretation of results due to concurrent changes in multiple parameters, including CAR sequence, antigen identity and expression level, and the biophysical characteristics of the CAR-antigen binding interaction (Frigault et al., 2015; Gomes-Silva et al., 2017; Long et al., 2015). Here, we began by examining the hypothesis that the amino-acid sequence of the ligand-binding domain of a CAR can significantly impact receptor activity, independent of the target-antigen identity or binding affinity. We chose to focus on CD20 CARs as our test platform because CD20 is a clinically validated antigen and enables direct benchmarking against the CD19 CAR in the treatment of B-cell lymphoma. A panel of CD20 CARs was constructed with single-chain variable fragments (scFvs) derived from four different monoclonal antibodies with similar K_D_ values (Mossner et al., 2010; Reff et al., 1994; Uchiyama et al., 2010) and similar complementarity determining region (CDR) structures predicted by abYsis (Martin and Thornton, 1996; Swindells et al., 2017). Three of the scFvs (Leu16, rituximab, and GA101) bind overlapping epitopes on the major extracellular loop of CD20, while ofatumumab binds both the minor and major loops of CD20 (Klein et al., 2013; Niederfellner et al., 2011; Rufener et al., 2016; Teeling et al., 2006) (Figure 1A; Figure S1A). All four CARs have identical extracellular spacer, transmembrane, and cytoplasmic signaling domains. This panel enables attribution of functional differences specifically to the scFv sequence while eliminating confounding factors such as differences in binding affinity and binding-epitope location. To better explore the potential relationship between scFv sequence and CAR tonic signaling, we chose to incorporate CD28 as the co-stimulatory domain, as CD28-containing CARs had previously been reported to be more prone to tonic signaling than 4-1BB– containing CARs (Frigault et al., 2015; Long et al., 2015).

**Figure 1.**
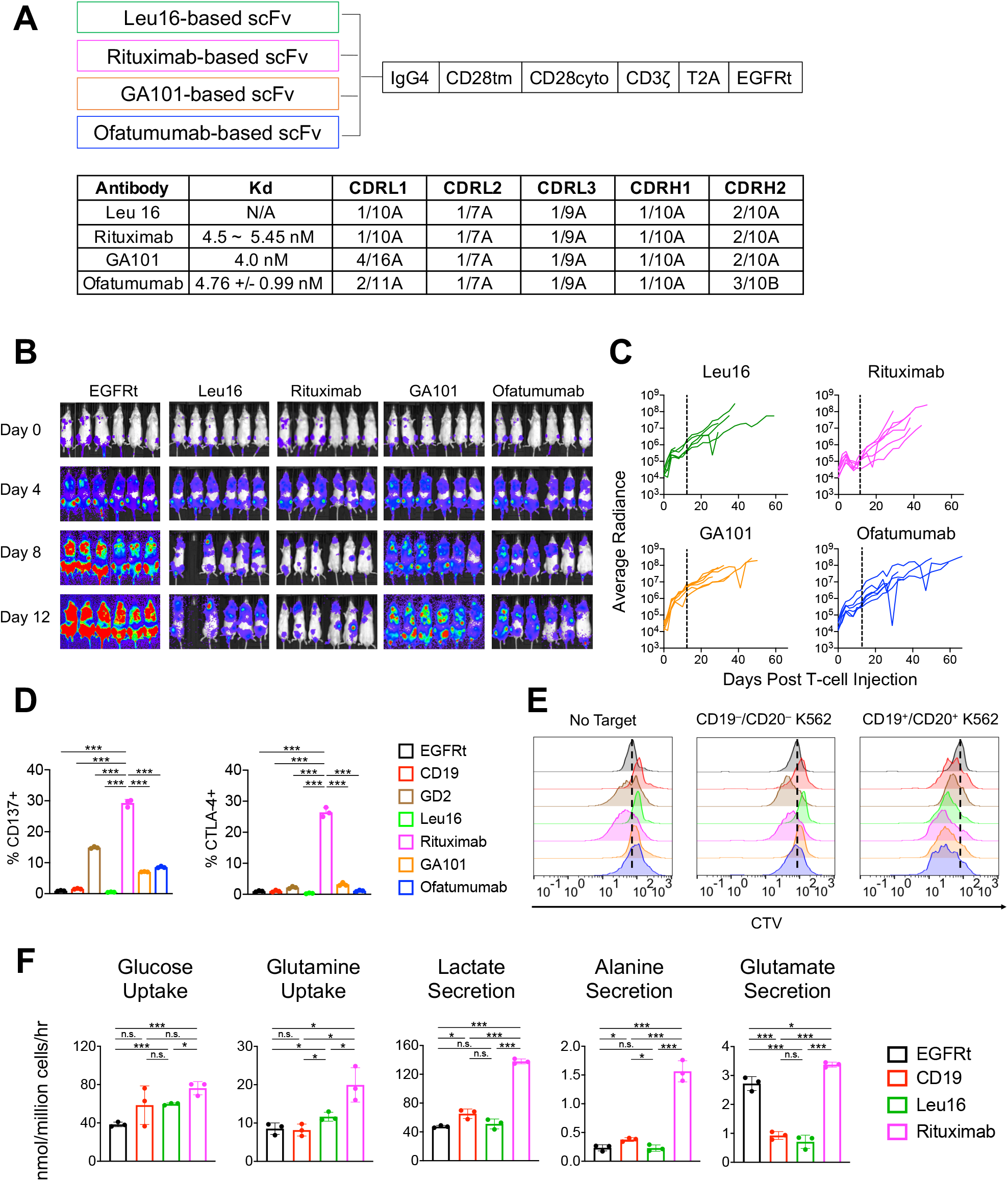
scFv sequence alters tonic signaling and CAR-T cell metabolism. (A) Schematic of a panel of 2^nd^-generation anti-CD20 CARs composed of scFvs derived from four different monoclonal antibodies fused to an IgG4 spacer, CD28 transmembrane and cytoplasmic domains, and CD3ζ signaling domain. The CAR is further fused via a self-cleaving T2A peptide to a truncated EGFR (EGFRt), which was used as transduction marker (top). K_D_ values and CDR structure-family designations of the four antibodies from which scFvs were derived (bottom). FR and CDR sequences and structure-family designations were determined as previously described (Chothia and Lesk, 1987; Chothia et al., 1989; Kabat and Wu, 1971; Martin and Thornton, 1996). (B–C) NSG mice were injected intravenously (i.v.) with 0.5 × 10^6^ firefly-luciferase–expressing Raji cells 6 days prior to treatment with 5 × 10^6^ CD8^+^ CD20-targeting CAR-T cells delivered i.v. (B) Tumor progression was monitored by bioluminescence imaging (n = 6 mice per group). Minimum and maximum values on the radiance-intensity scale are 5 × 10^4^ and 1 × 10^7^, respectively. (C) Average radiance (p/sec/cm2/sr) of individual animals for each test group. Black dotted line denotes day 12 post T-cell injection, the time at which the rituximab CAR-T cell group began to rapidly lose tumor control. (D) Activation and exhaustion maker expression on CAR^+^ T cells were evaluated 11 days post Dynabead removal, without CD20 antigen stimulation. Data bars indicate the means of technical triplicates ± 1 standard deviation (S.D.). Results are representative of three independent experiments using T-cells derived from three different healthy donors. Unless otherwise noted, *p* values were determined by unpaired, two-tailed, two-sample Student’s t-test; * *p*<0.05, ** *p*<0.01, *** *p*<0.001, n.s. not statistically significant. (E) A 4-day T-cell proliferation assay on CAR^+^ T cells with CellTrace Violet (CTV) dye in the absence or presence of target cells (on-target, CD19^+^/CD20^+^ K562 cells; off-target, parental K562 cells) at 2:1 effector-to-target (E:T) ratio. Data shown are representative of five independent experiments from five different healthy donors. (F) Metabolic rates of CD20 CAR-T cells cultured for 24 hours in RPMI supplemented with 10% dFBS and exogenous IL-2 and IL-15, without CD20 antigen stimulation. Data bars indicate the means of technical triplicates ± 1 S.D. Results are representative of three independent experiments from three different healthy donors. \**p*<0.05, \*\**p*<0.01, \*\*\**p*<0.001, n.s. not statistically significant.

All four CD20 CAR variants were efficiently expressed on the surface of primary human T cells (Figure S1B), and T cells expressing each CAR grew at comparable rates during *ex vivo* expansion with cytokine support (Figure S1C). In response to repeated antigen stimulation, all CAR-T cell variants demonstrated the ability to lyse target cells and proliferate (Figure S1D). However, despite similar performances during *in vitro* functional assays, T cells expressing the four CD20 CAR variants showed distinct *in vivo* tumor-killing dynamics in the Raji xenograft model (Figure 1B). In particular, CD8^+^ rituximab-based CAR-T cells exerted substantially stronger tumor control at early time points compared to all other CD20 CAR-T cells tested (Figure 1B). And yet, starting approximately two weeks post T-cell injection, animals treated with rituximab CAR-T cells began exhibiting accelerated tumor growth—faster than any other test group (Figure 1C; Figure S2A)—suggesting rapid onset of functional exhaustion among the rituximab CAR-T cells.

Animals in all test groups carried detectable populations of CAR-T cells at the time of sacrifice, particularly in the liver, spleen, and tumor metastases recovered from the brain (Figure S2B,C). However, nearly all T cells were PD-1+ and many showed elevated LAG-3 expression (Figure S2D,E). Combined with the failure to eradicate tumor, these results suggest all four groups of CAR-T cells were dysfunctional by the end of the study, despite their persistence *in vivo*.

We next sought to elucidate what biological differences among the four CD20 CAR-T cell variants could explain the different temporal dynamics of their *in vivo* tumor response, and whether the underlying biology could shed light on improved CAR designs capable of sustained tumor control. In particular, we wished to understand why the rituximab CAR led to initially robust but ultimately short-lived anti-tumor response, and whether we could prolong anti-tumor efficacy through rational CAR protein engineering.

In the absence of antigen stimulation, T cells expressing the rituximab CAR showed clear activation-marker expression (Figure 1D) and cell proliferation (Figure 1E), to a similar or stronger degree than T cells expressing the 14g2a-based GD2 CAR that is known to tonically signal (Long et al., 2015). These observations indicated clear basal activation of rituximab CAR-T cells that was directly attributable to the CAR construct, and led us to hypothesize that rituximab CAR-T cells could rapidly initiate anti-tumor response in part due to this pre-inclination toward T-cell activation. It has been shown that initiation of robust T-cell effector function requires support from aerobic glycolysis (Chang et al., 2013). Therefore, to examine the hypothesis that rituximab CAR-T cells are potentiated toward rapid effector function, we set out to characterize its basal metabolic state by measuring the rates of nutrient uptake and byproduct secretion. In the absence of antigen stimulation, rituximab CAR-T cells exhibited significantly faster glucose, glutamine, and amino acid uptake compared to mock-transduced T cells and T cells expressing either the CD19 CAR or the Leu16-based CD20 CAR (Figure 1F; Figure S3). Furthermore, rituximab CAR-T cells showed 2–7-fold increases in lactate and alanine secretion, as well as >3-fold increase in glutamate secretion compared to the other CAR-T cells, indicating dramatically increased glycolytic flux and glutaminolysis, respectively. As glucose and glutamine are responsible for the majority of ATP production, faster glycolysis and glutaminolysis imply greater energy expenditure, consistent with the behavior of activated T cells (Chang et al., 2013).

Taken together, these observations indicate that, in the absence of antigen stimulation, rituximab CAR-T cells are basally activated and metabolically equipped to initiate T-cell effector function. This heighted metabolism at rest may explain rituximab CAR-T cells’ rapid anti-tumor activity at early time points *in vivo*, but chronic, low-level T-cell activation in the absence of antigen stimulation may also have contributed to rituximab CAR-T cells’ accelerated loss of tumor control compared to the other CAR-T cell variants, as observed in our animal study. To evaluate this possibility and its implication on CAR design, we next investigated whether tuning the level of basal activation could yield CAR-T cells that are capable of both rapid and sustained anti-tumor response.

### Torsional reorientation of signaling domain tunes CAR-T cell activity

Receptor dimerization at the cell surface can trigger CAR signaling (Chang et al., 2018), and an earlier study suggested that tonic signaling may be a consequence of antigen-independent CAR clustering caused by certain scFv sequences (Long et al., 2015). We verified that all four CD20 CAR variants shown in Fig. 1A are uniformly distributed on the T-cell surface in the absence of antigen stimulation (Figure S4A), thus ruling out macro-scale CAR clustering as the cause of tonic signaling by the rituximab-based CAR. Nevertheless, the signaling cascade downstream of CD3ζ involves adaptor proteins and kinases whose interactions with the receptor chain are directly impacted by the physical proximity and conformation of receptor chains (Hartman and Groves, 2011). We thus hypothesized that tonically signaling CARs may adopt a conformation that enables basal receptor signaling, and that altering the conformation of the CAR molecule could enable the tuning of CAR signaling intensity both in the absence and in the presence of antigen stimulation, thereby impacting downstream CAR-T cell behavior.

To systematically investigate the effect of CAR-conformation change, we inserted one to four alanines between the transmembrane and cytoplasmic domains of CD28 in the rituximab-based CAR, with each alanine expected to cause a ∼109° turn in the protein structure through the formation of an alpha helix (Constantinescu et al., 2001; Liu et al., 2008; Scheller et al., 2018) (Figure 2A; Figure S5A). This “twisting” process is designed to alter the alignment between the CAR’s extracellular ligand-binding domain and its cytoplasmic signaling domains, potentially impacting the packing behavior of receptors in high-density areas (e.g., within microclusters upon antigen-ligation), as well as the accessibility of docking and phosphorylation residues involved in downstream signaling.

**Figure 2.**
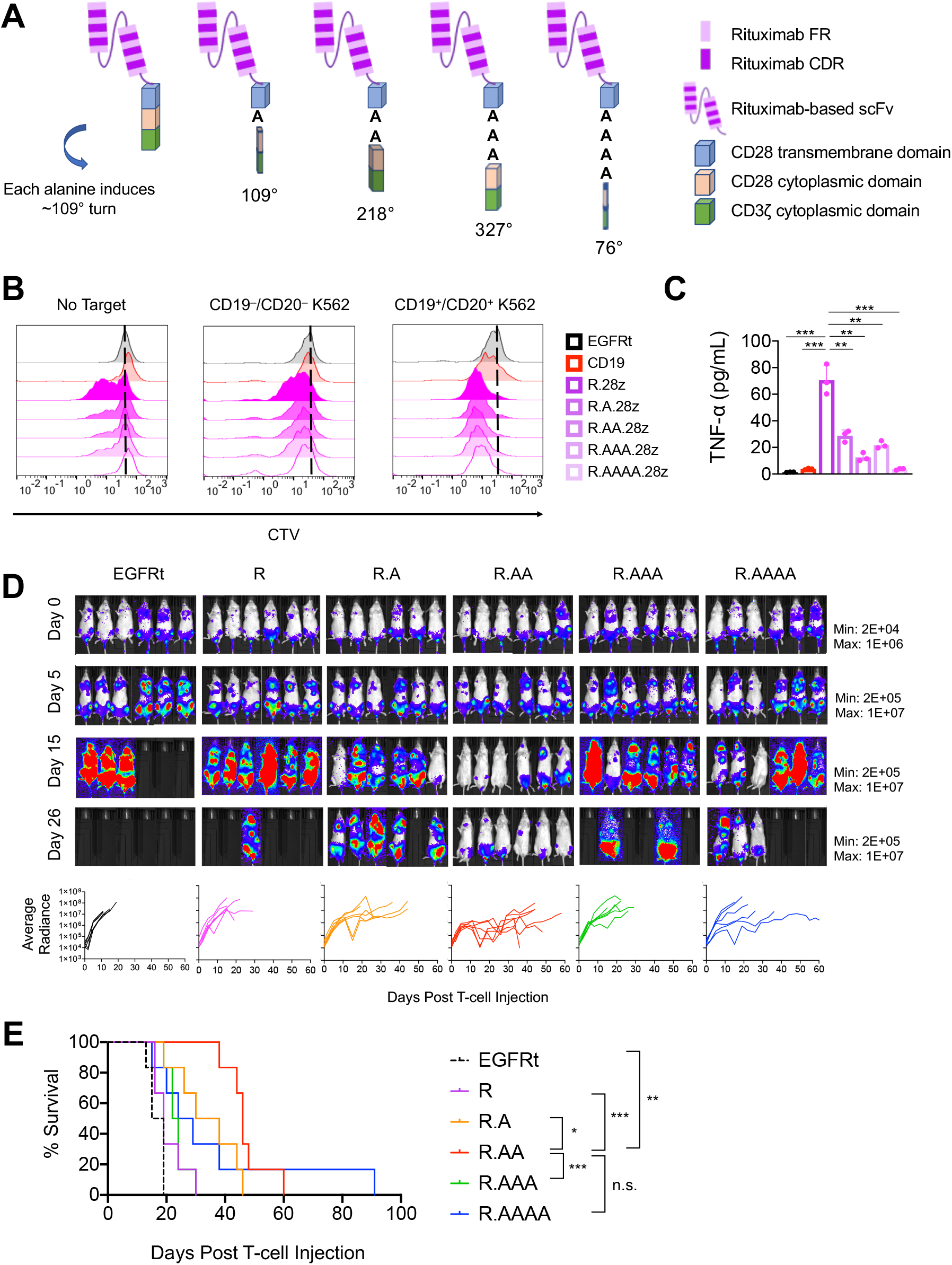
Torsional reorientation of signaling domain tunes CAR-T cell activity. (A) Schematic of alanine incorporation into the Rituximab-based CAR. (B) A 4-day T-cell proliferation assay with CellTrace Violet (CTV) dye in the absence or presence of target cells at 2:1 E:T ratio. Data shown are representative of three independent experiments from three different healthy donors. (C) TNF-α production of CAR-T cells was measured 7 days post Dynabead removal, without CD20 antigen stimulation. Results are representative of three independent experiments from three different healthy donors. \**p*<0.05, \*\**p*<0.01, \*\*\**p*<0.001, n.s. not statistically significant. (D,E) NSG mice were injected intravenously with 0.5 × 10^6^ firefly-luciferase–expressing Raji cells followed by two doses of CAR^+^ T cells 6 days (1.35 × 10^6^ cells) and 12 days (1.5 × 10^6^ cells) later; n = 6 mice per group. This and all subsequent *in vivo* studies utilized T cells generated from a naïve/memory (CD14^−^/CD25^−^/CD62L^+^ sorted) starting population (see Materials and Methods section for discussion on T-cell subtype usage). (D) Tumor progression was monitored by bioluminescence imaging (top). Average radiance (p/sec/cm2/sr) of individual animals are shown for each group (bottom). (E) Kaplan-Meier survival curve. Statistical significance was determined by log-rank (Mantel-Cox) test. \**p*<0.05, \*\**p* 0.01, \*\*\**p*<0.001, n.s. not statistically significant.

All alanine-insertion variants as well as the original rituximab CAR expressed well on the T-cell surface (Figure S5B), demonstrated similar ability to lyse target cells and proliferate in response to repeated antigen challenge (Figure S5C), and showed similar levels of activation and exhaustion marker expression in the absence of antigen stimulation (Figure S5D). However, alanine insertion resulted in a clear reduction of antigen-independent T-cell proliferation and TNF-α production compared to the original rituximab-based CAR (Figure 2B,C), indicating the insertion of torsional linkers significantly reduced basal CAR-T cell activity. The impact of CAR structural alteration was evident *in vivo*, with T cells expressing rituximab-based CARs containing one, two, or four alanines showing significantly improved control of Raji xenografts compared to the original rituximab CAR (Figure 2D). In particular, the two-alanine CAR increased median survival period by 2.1-fold compared to the original rituximab CAR (55 days vs. 26 days; Figure 2E).

In principle, the insertion of alanine-based torsional linkers can be performed on any CAR protein, irrespective of the particular ligand-binding or signaling domains incorporated in the CAR. To probe the generalizability of this CAR-tuning method, we evaluated the effects of alanine insertion in the 14g2a-based GD2 CAR. All five GD2 CAR variants (with zero to four alanines inserted between the transmembrane and cytoplasmic domains) expressed well on the T-cell surface, and exhibited similar anti-tumor responses upon repeated antigen challenge *in vitro* (Figure S6A–C). Consistent with results from the CD20 CAR panel (Figure 2D), T cells expressing GD2 CARs containing one, two, or four alanines outperformed GD2 CARs with zero or three alanine inserted (Figure S6D). Furthermore, analysis of liver and spleen recovered at the time of sacrifice revealed a significantly higher level of T cells expressing the two-alanine CAR construct, underscoring the utility of two-alanine insertion in CARs containing CD28 transmembrane and cytoplasmic domains (Figure S6E).

Taken together, these results indicate that alanine insertion into CAR molecules is a generalizable method to tune CAR-T cell activation and strengthen anti-tumor efficacy *in vivo*. However, despite the improvements seen with alanine insertion, the CD20 CAR-T cells remained unable to eradicate Raji xenografts in our mouse model (Figure 2D,E). We thus set out to explore additional design parameters that could further enhance CAR-T cell function.

### scFv sequence hybridization yields superior CAR variant

The observation that the rituximab CAR’s behavior was distinct among the original panel of four CD20 CARs was striking given that all CARs had identical components aside from the scFv (Figure 1A). Furthermore, the rituximab scFv has 91% sequence identity compared to the Leu16 scFv (Figure S1A), yet Leu16 and rituximab CAR-T cells had dramatically different behaviors in terms of both basal T-cell activation and *in vivo* tumor-killing dynamics (Figure 1B– E). These results indicate that limited variations in scFv sequence can exert a significant impact on CAR signaling activities.

Each scFv comprises a light chain and a heavy chain, and each chain can be further subdivided into four framework regions (FRs) flanking three complementarity-determining regions (CDRs) (Figure 2A). It has previously been suggested that the FRs of certain scFvs such as 14g2a may be the cause of tonic signaling and CAR-T cell exhaustion, and it was demonstrated that incorporation of the 14g2a scFv’s FRs into the CD19 CAR resulted in a hybrid receptor that triggered T-cell exhaustion (Long et al., 2015). Thus it was possible to impair a CAR through scFv alterations, but it remained unclear whether a CAR can be improved through scFv sequence hybridization. Our results thus far indicated that lowering the intensity of CAR tonic signaling via alanine insertion can improve CAR-T cell function. We next explored whether changes in scFv through sequence hybridization could provide a second “knob” by which CAR signaling activities could be rationally tuned.

Leu16 and rituximab were chosen as the starting point for hybridization due to their similar sequences yet disparate activity profiles. We constructed hybrid CARs whose scFv comprised the FRs of rituximab and CDRs of Leu16 (RFR-LCDR), or vice versa (LFR-RCDR) (Figure 3A). The RFR-LCDR hybrid CAR triggered robust tumor-cell lysis and T-cell proliferation upon repeated antigen challenge, providing the first example of a CD20 CAR whose functionality is comparable to that of the CD19 CAR in our *in vitro* assays (Figure 3B). RFR-LCDR hybrid CAR-T cells showed clear activation marker expression in the absence of antigen stimulation, indicating a non-zero level of T-cell activation at rest (Figure 3C). In contrast, the LFR-RCDR hybrid was completely non-functional despite being efficiently expressed on the T-cell surface (Figure 3B; Figure S7A), thus it was excluded from further characterization. Henceforth, the term “hybrid CAR” refers to the RFR-LCDR variant.

**Figure 3.**
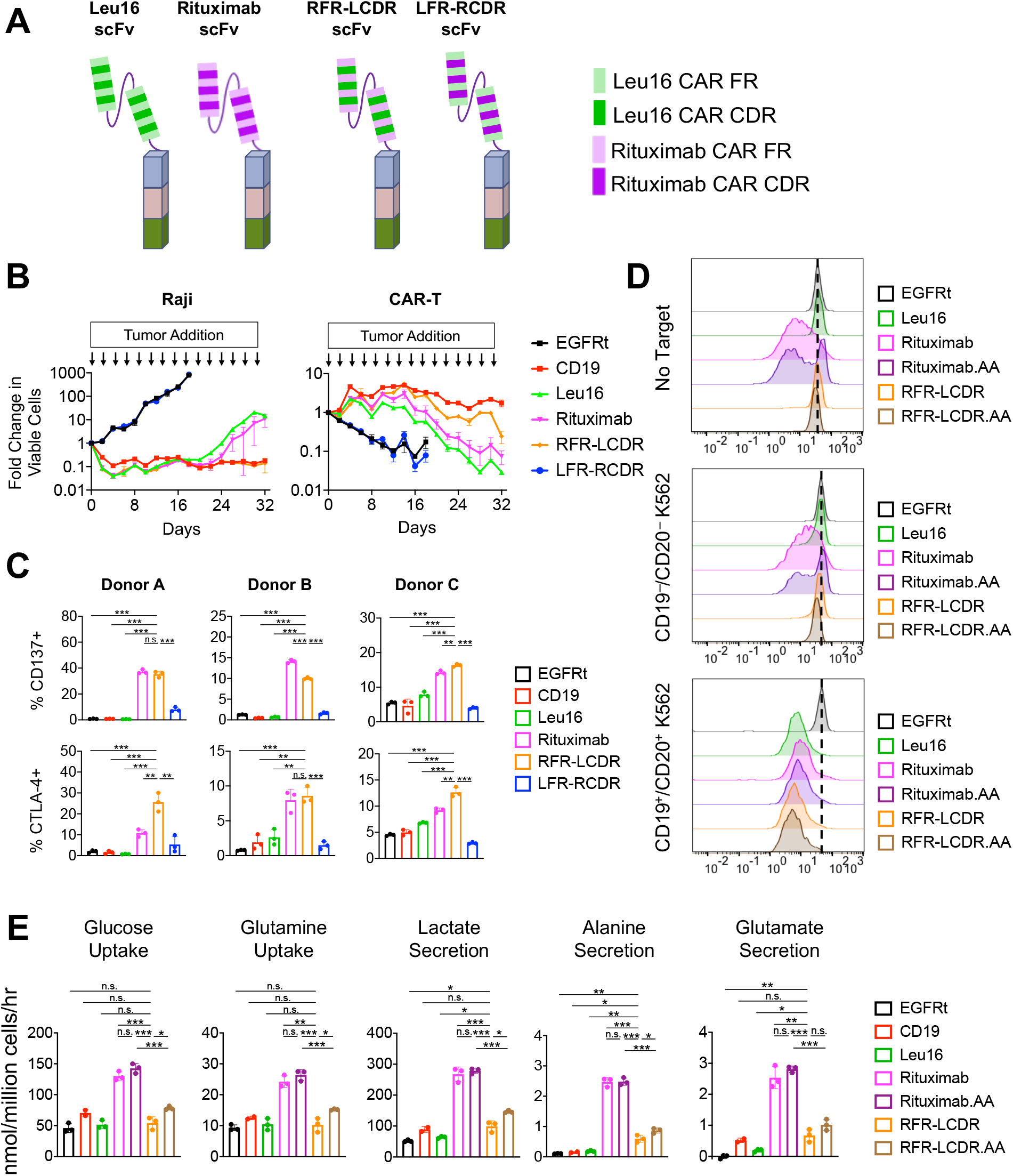
scFv sequence hybridization yields functionally superior CAR variant. (A) Schematic of scFv sequence hybridization in CAR molecules. The Framework regions (FR) and complementarity-determining regions (CDRs) of Leu16- and rituximab-derived scFvs were intermixed to yield two new CAR variants. (B) RFR-LCDR hybrid CAR-T cells show superior anti-tumor function and T-cell proliferation upon repeated antigen challenge. CAR-T cells were challenged with Raji cells (CD19^+^/CD20^+^) at a 2:1 E:T ratio every two days. T-cell and target-cell counts were quantified by flow cytometry. Data shown are the means of technical triplicates ± 1 S.D. Results are representative of three independent experiments from three different healthy donors. (C) Activation and exhaustion maker expression was evaluated 11 days post Dynabead removal, without CD20 antigen stimulation. Data bars indicate the means of technical triplicates ± 1 S.D. Results are representative of three independent experiments from three different healthy donors. \**p*<0.05, \*\**p*<0.01, \*\*\**p*<0.001, n.s. not statistically significant. (D) A 4-day CAR-T cell proliferation assay with CellTrace Violet (CTV) dye in the absence or presence of target cells at 2:1 E:T ratio. (E) Metabolic analysis of CAR-T cells in culture in the absence of antigen stimulation. CAR-T cells were cultured for 72 hours in RPMI supplemented with 10% heat-inactivated, dialyzed fetal bovine serum (HI-dFBS), IL-2, and IL-15. Data bars indicate the means of technical triplicates ± 1 S.D. Data are representative of three independent experiments from three different healthy donors. \**p*<0.05, \*\**p*<0.01, \*\*\**p*<0.001, n.s. not statistically significant.

A modified hybrid CAR with two alanines inserted between the transmembrane and cytoplasmic CD28 signaling domains (RFR-LCDR.AA) was included in subsequent studies to determine whether sequence hybridization and alanine insertion would synergize to further improve CAR-T cell function. Both hybrid CARs showed similar surface distribution as rituximab-based CARs with and without alanine (Figure S4), and all CAR-T cells produced Th1 cytokines in response to antigen stimulation (Figure S7B), confirming tumor reactivity. However, hybrid CAR-T cells showed no sign of antigen-independent T-cell proliferation (Figure 3D), and scFv sequence hybridization significantly reduced TNF-α production and increased IL-2 production in the absence of antigen stimulation compared to rituximab-based CAR-T cells (Figure S7C). Furthermore, metabolic analysis on CAR-T cell culture supernatant revealed that scFv hybridization significantly reduced glucose and glutamine uptake as well as glutamate, alanine, and lactate secretion in the absence of antigen stimulation (Figure 3E; Figure S8). Consequently, the metabolic rates for hybrid CAR-T cells at rest were at an intermediate level, between those of T cells expressing either of the two parental constructs (i.e., rituximab or Leu16 CARs). Taken together, these results indicate scFv hybridization can significantly reduce the intensity of tonic signaling without completely abolishing basal T-cell activation.

We next performed head-to-head comparisons of the hybrid CAR against each of its parent constructs and the CD19 CAR *in vivo* (Figure 4A). Results indicate that RFR-LCDR CAR-T cells efficiently rejected both the original tumor as well as tumor re-challenge (Figure 4B,C). In fact, this marked the first example in our experience of a CD20 CAR that outperforms the CD19 CAR in eradicating B-cell lymphoma xenografts.

**Figure 4.**
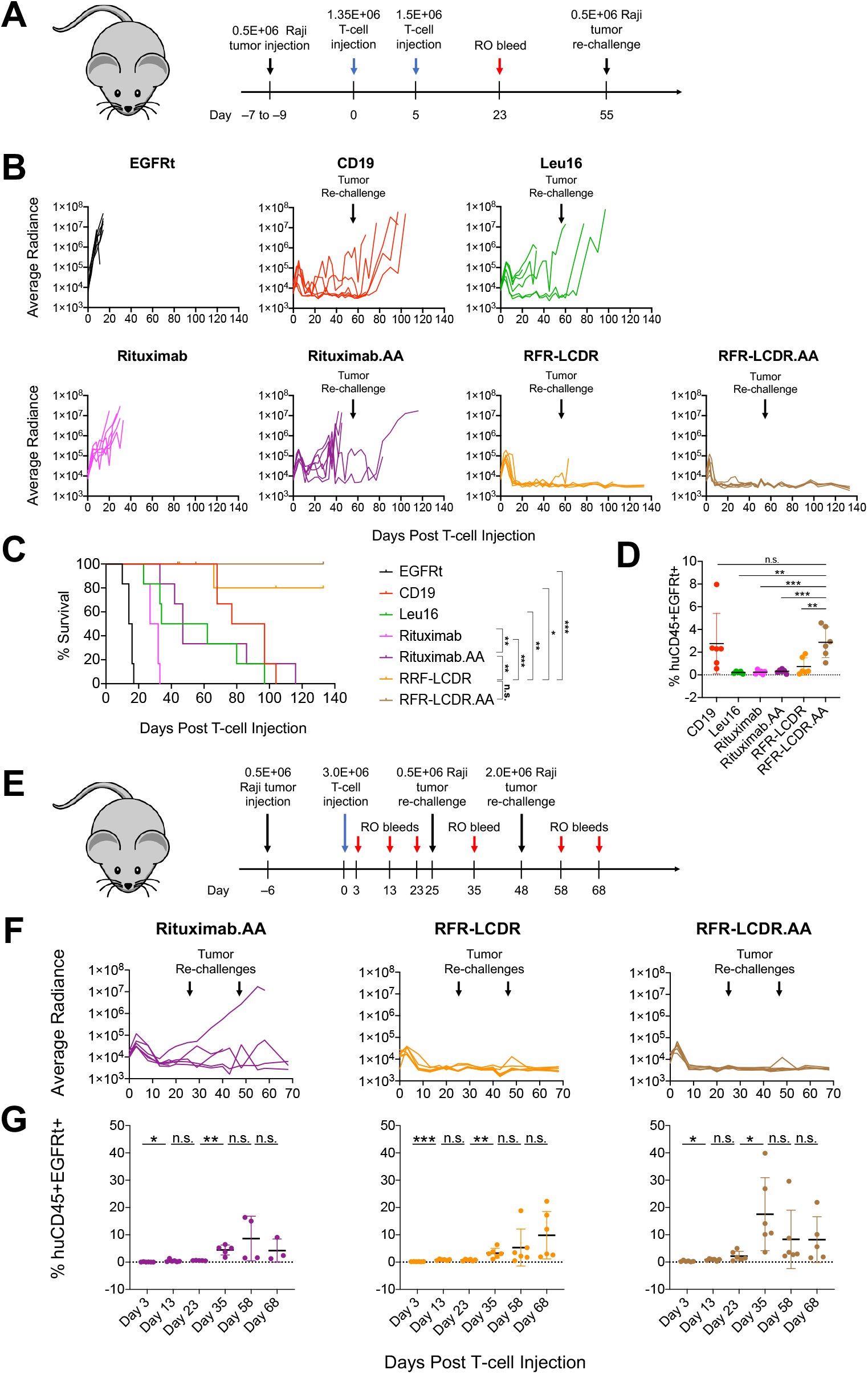
scFv hybridization in combination with torsional reorientation in CAR protein further enhance CAR-T cell function *in vivo*. (A–D) NSG mice were injected intravenously with firefly-luciferase–expressing Raji cells followed by two doses of CAR^+^ T cells, and one Raji tumor rechallenge. (A) Schematic of *in vivo* experiment (n = 6 mice per group). (B) Tumor signal in individual animals quantified by bioluminescence imaging. (C) Kaplan-Meier survival curve. Log-rank (Mantel-Cox) test was performed for pair-wise comparisons. \**p*<0.05, \*\**p*<0.01, \*\*\**p*<0.001, n.s. not statistically significant. (D) Frequency of human CD45^+^EGFRt^+^ cell in peripherical blood collected from mice at Day 23 after first dose of T-cell infusion. (E–G) NSG mice were injected intravenously with firefly-luciferase expressing Raji cells followed by a single dose of CAR^+^ T cells. Mice were re-challenged twice with Raji cells, on Day 25 and Day 45 post T-cell injection. (E) Schematic of *in vivo* experiment (n = 6 mice per group). (F) Tumor signal in individual animals quantified by bioluminescence imaging. (G) Frequency of human CD45^+^EGFRt^+^ cell in peripherical blood collected from mice over time. \**p*<0.05, \*\**p*<0.01, \*\*\**p*<0.001, n.s. not statistically significant.

Consistent with prior results, alanine insertion substantially improved the rituximab CAR (Figure 4B,C). Since RFR-LCDR and RFR-LCDR.AA CAR-T cells both conferred nearly perfect protection against tumor and re-challenge, we could not assess whether alanine insertion in the hybrid CAR context provided further benefits in tumor killing. However, RO blood analysis indicated that RFR-LCDR.AA CAR-T cells had superior *in vivo* survival and expansion compared to RFR-LCDR CAR-T cells (Figure 4D). These results were further validated in a second *in vivo* study where animals were treated with a single (instead of split) dose of CAR-T cells and re-challenged with tumor cells twice, with escalating tumor dosage levels (Figure 4E,F). Rituximab.AA, RFR-LCDR, and RFR-LCDR.AA CAR-T cells all persisted and rebounded after each tumor re-challenge, with RFR-LCDR.AA CAR-T cells showing the highest level of CAR-T cells in peripheral blood both before and after the first tumor re-challenge (Figure 4G). Both hybrid CAR variants conferred complete protection to the animals through the study.

Taken together, these results indicate that the functionality of a CAR can be improved by changing its scFv sequence and that tuning—rather that complete elimination—of basal T-cell activation could result in CAR designs with significantly improved anti-tumor functions.

### Memory phenotype enrichment and minimization of CAR-driven metabolic disturbance enhance CAR-T cell function

To better understand the biology that underpins the functional differences observed *in vivo*, we performed RNA-seq and ATAC-seq analyses on rituximab, Leu16, RFR-LCDR, and RFR-LCDR.AA CAR-T cells recovered from tumor-bearing animals 9 days post T-cell injection (Figure S9). Results revealed stark differences in the transcriptomic and epigenetic profiles of rituximab CAR-T cells compared to the others, with the RFR-LCDR.AA CAR being the most distant from the rituximab CAR (Figure 5A,B; Figure S10). Compared to each of the other CAR variants, rituximab CAR-T cells showed significantly increased expression of *ATP9A*, which encodes for a phospholipid flippase that positively regulates GLUT1 recycling (Tanaka et al., 2016) (Figure 5A,B). GLUT1 recycling from endosomes to the plasma membrane is essential for cellular glucose uptake in support of glycolysis, and different GLUT1 expression levels have been shown to correlate with distinct effector functions in human T cells (Cretenet et al., 2016; Macintyre et al., 2014). These results echoed our previous observation that rituximab and rituximab.AA CAR-T cells exhibit elevated glucose uptake at rest (Figure 3E), and prompted us to analyze the glucose level in blood-serum samples that had been collected from the *in vivo* study shown in Figures 4E–G. The results revealed significantly lower glucose levels in the serum of mice treated with rituximab.AA CAR-T cells (Figure 5C), further supporting the notion that rituximab-based CAR-T cells are substantially more glycolytically active than hybrid CAR-T cells.

**Figure 5.**
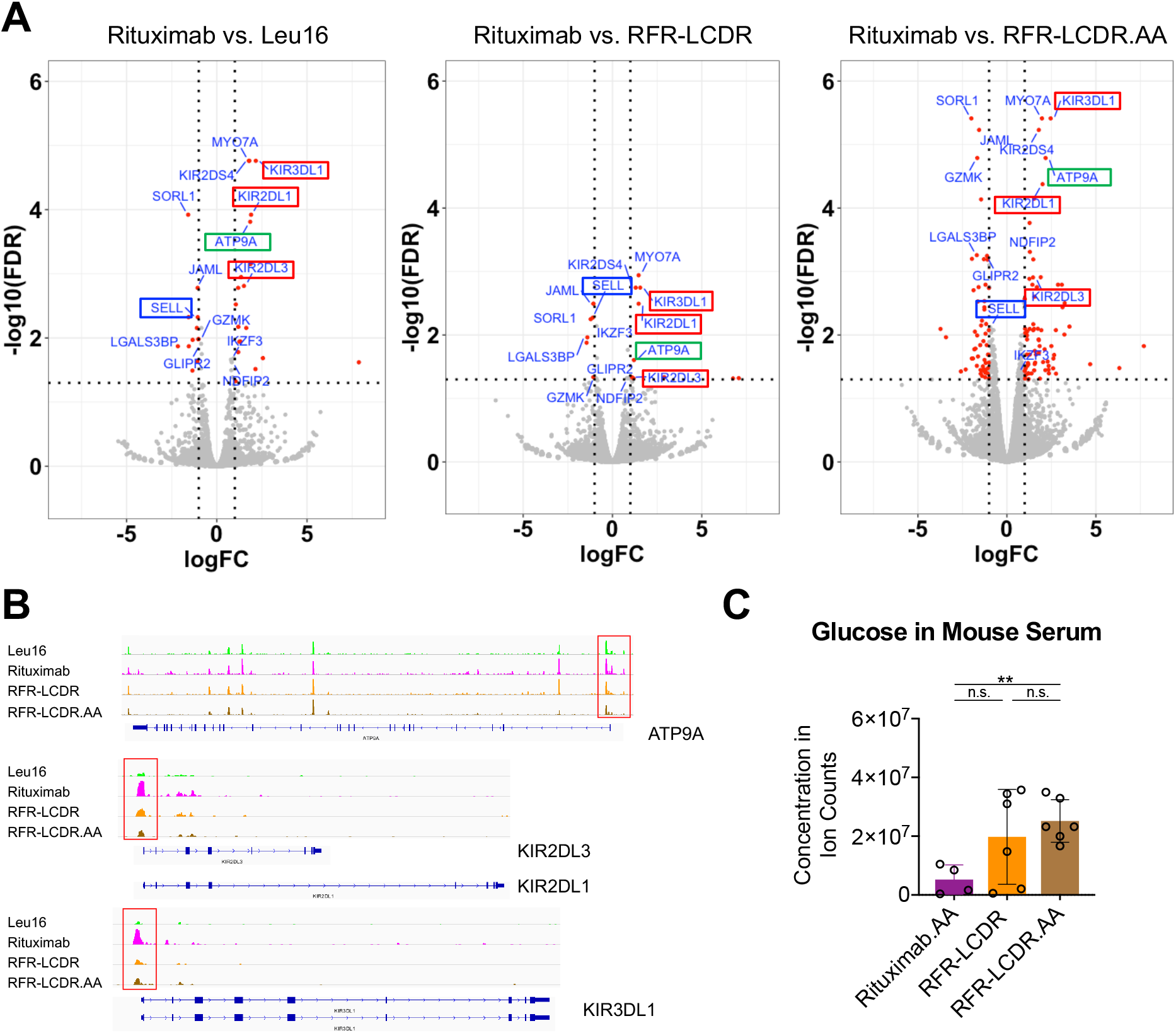
Transcriptomic and epigenetic analyses reveal CAR-dependent variations in T-cell phenotypes. (A,B) NSG mice were injected i.v. with 0.5 × 10^6^ firefly-luciferase–expressing Raji cells 6 days prior to treatment with 2.85 × 10^6^ CAR^+^ T cells delivered i.v. Liver, spleen, cardiac blood, and bone marrow were collected from tumor-bearing mice 9 days after T-cell injection (n = 2 mice per group). CAR^+^ T cells were obtained by enriching for huCD45^+^ EGFRt^+^ populations, and subsequently analyzed by RNA-seq and ATAC-seq. (A) Volcano plots of differentially expressed genes of Rituximab CAR-T cells versus Leu16 (left), RFR-LCDR (middle), or RFR-LCDR.AA (right) CAR-T cells based on RNA-seq. All differentially expressed genes are plotted in grey. Genes that have at least 2-fold upregulation or downregulation (log_2_FC > 1 or log_2_FC < −1) with FDR < 0.05 are shown as red dots. The names of genes that appear in all three sets of pair-wise comparisons with log_2_FC > 1 and FDR < 0.05 or log_2_FC < −1 and FDR < 0.05 are labeled. (B) Genome browser files of differentially accessible regions in Leu16, Rituximab, RFR-LCDR, RFR-LCDR.AA CAR-T cells at the *ATP9A, KIR2DL3, KIR2DL1, KIR3DL1* loci. (C) Animals treated with Rituximab-based CAR-T cells show reduced glucose levels in serum. Blood serum was collected from animals in the study shown in Figure 4E–G on Day 58 post T-cell injection. Glucose concentration in serum was measured by LC-MS. Data bars indicate the means of biological replicates ± 1 S.D. (n = 6 for both hybrid CAR-T cell groups; n = 4 for Rituximab.AA CAR-T cell treatment group due to death caused by tumor burden prior to sample collection date). \**p*<0.05, \*\**p*<0.01, \*\*\**p*<0.001, n.s. not statistically significant.

In addition to increased glycolysis, rituximab CAR-T cells recovered from mice showed significantly lower CD62L expression (encoded by the *SELL* gene) and higher transcript levels for the inhibitory receptors KIR2DL1, KIR2DL3, and KIR3DL1 (Bjorkstrom et al., 2012) compared to each of the other three CAR-T cell types (Figure 5A). Upregulation of the inhibitory KIRs was further confirmed by ATAC-seq (Figure 5B). Taken together, rituximab CAR-T cells exhibit a phenotype consistent with effector T cells trending toward functional exhaustion.

In contrast, hybrid CAR-T cells, particularly RFR-LCDR.AA CAR-T cells, are significantly enriched in the memory phenotype based on Gene Set Enrichment Analysis (GSEA) (Figure 6A,B). GSEA also revealed that both hybrid CAR-T cell populations recovered from mice exhibit significantly stronger T-cell activation, cytotoxicity, receptor-signaling, and interferon gamma signatures compared to rituximab and Leu16 CAR-T cells (Figure 6A,C). For RFR-LCDR.AA CAR-T cells in particular, this strong T-cell activation signature coexists with low cell-cycle and DNA-replication activity, consistent with the memory phenotype and in stark contrast with rituximab CAR-T cells (Figure 6A,D). Of note, the enrichment of memory subtype among RFR-LCDR.AA CAR-T cells was not evident prior to injection into tumor-bearing mice (Figure S11), indicating the change occurred upon encountering antigen stimulation and/or the *in vivo* environment.

**Figure 6.**
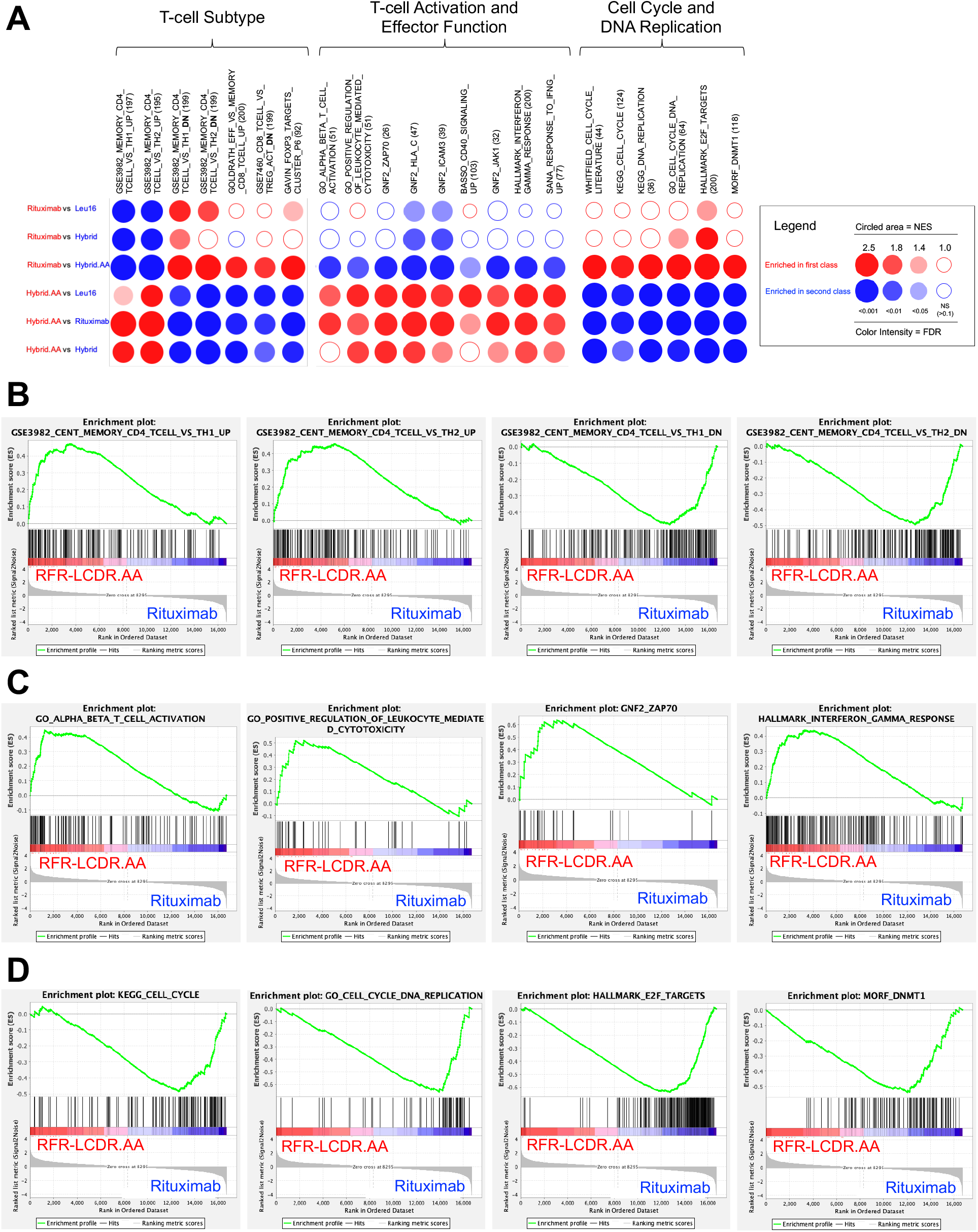
RFR-LCDR.AA CAR-T cells show robust T-cell activation coupled with memory phenotype. (A) Gene Set Enrichment Analysis (GSEA) was performed on RNA-seq data obtained as described in Figure 5A. A summary of results in pathways related to T-cell phenotype and function are shown in BubbleGUM map format (Spinelli et al., 2015). (B) Mountain plots of pathways related to T-cell subtype. (C) Mountain plots of pathways related to T-cell activation and interferon gamma signaling. (D) Mountain plots of pathways related to cell cycle, DNA replication, and cell metabolism.

Taken together, our data indicate that rituximab CAR-T cells exhibit elevated glycolysis both at rest and upon antigen exposure *in vivo*. In contrast, RFR-LCDR.AA, a novel CAR obtained through scFv sequence hybridization combined synergistically with alanine insertion, promotes the formation of memory T cells that can mount robust effector functions while maintaining relatively low metabolic activity levels as well as long-term persistence *in vivo*.

## DISCUSSION

In this study, we explored the effect of rational CAR sequence and structural modifications on the behavior of CAR-T cells, and observed that even modest alterations—as small as the insertion of two alanine residues—can induce dramatic changes in the anti-tumor efficacy of the resulting CAR-T cells. We found tonic signaling—and the associated basal T-cell activation—to be a useful measure by which to quantify and tune CAR behavior. Our approach of engineering CARs to exhibit a low but non-zero levels of tonic signaling yielded CAR-T cells that execute rapid response to antigen stimulation while enabling sustained anti-tumor activity. Methods such as the insertion of torsional linkers comprised of alanines and the hybridization of FR and CDR sequences from different scFvs enable the generation of CAR variants that fall along the spectrum of tonic-signaling intensities, and some of these variants could outperform even the gold-standard CD19 CAR in *in vivo* tumor control.

Alanine insertion is a generalizable method for tuning CAR tonic signaling as well as antigen-stimulated CAR-T cell function. In this study, we found the insertion of two alanines between the transmembrane and cytoplasmic domains of CD28 to improve the *in vivo* function of both CD20 and GD2 CARs. Given that the intended effect of alanine insertion is to alter or “twist” the CAR conformation, and the starting protein conformation is likely dependent on the specific components incorporated in the CAR, the optimal number of alanine may vary with the specific transmembrane and cytoplasmic domains used in the CAR. Detailed biochemical analysis of the downstream signal-transduction pathway could further elucidate the molecular mechanism that underpins the functional improvements in our study, and inform on whether the alanine-insertion strategy could be adapted to CARs with a variety of transmembrane and cytoplasmic domains.

In addition to alanine insertion, scFv hybridization proved to be a fruitful method by which to generate improved CAR variants. In the case of the RFR-LCDR hybrid characterized in this study, a CAR built with FRs from a tonically signaling CAR (rituximab) and CDRs from a non-tonically signaling CAR (Leu16) led to a chimera with intermediate tonic-signaling intensity, and far-superior anti-tumor efficacy compared to either parent. This result highlights the unpredictability of the effect of scFv sequence on CAR-T cell function. In-depth analysis of protein structure and site-specific mutations may allow for precise identification of the truly critical residues and enable a fully rational approach to scFv design in the future.

In the course of this study, we used two different starting T-cell populations to evaluate the effect of CAR engineering, reflecting updates in our understanding of the most effective T-cell subtype to use for therapeutic applications. Our first *in vivo* study (Figure 1B,C) was performed using CD8^+^ T cells based on early literature indicating CD8^+^ cytotoxic T lymphocytes (CTLs) are a therapeutically promising cell type (Yee et al., 2002; Klebanoff et al., 2005; Powell et al., 2005). More recently, studies by our group and others suggest that naïve/memory T (T_N/M_) cells enriched for CD14^−^/CD25^−^/CD62L^+^ phenotype could yield greater T-cell function and may have increased clinical potential compared to CD8^+^ CTLs (NCT04007029; Zah et al., 2020; REF). To maximize the clinical relevance of our studies, we transitioned to using T_N/M_ cells in our later experiments, including those shown in Figures 2 and 4. This change in T-cell subtype, and associated changes in the T-cell dosages used, may explain the different *in vivo* anti-tumor dynamics observed in Figure 1 versus later studies. Consistent results from repeated comparisons of rituximab vs. rituximab.AA (Figures 2D and 4B), as well as rituximab.AA vs. hybrid vs. hybrid.AA (Figures 4B and 4F), provide reproducible evidence of the positive impacts of alanine insertion and scFv hybridization on *in vivo* anti-tumor efficacy.

In this work, tonic signaling is used as a guide to CAR protein engineering, as it provides a quantifiable phenotype where a certain level of tonic-signaling correlates with robust CAR-T cell function. However, the improved functional outcome is likely the result of not just tonic signaling itself, but a variety of additional, associated factors that changed with tonic signaling as we modified the CAR design. For example, a phenotype found to be strongly associated with tonic signaling—and tunable by the protein-engineering methods discussed above—is elevated metabolic activity at rest, particularly glycolysis and glutaminolysis. A significant body of literature has shown that increased glycolytic activity is characteristic of effector T cells, and that heightened glycolysis is essential to robust T-cell effector function (Bantug et al., 2018). The fact that rituximab CAR-T cells exhibit strong glycolysis both at rest and upon antigen stimulation is consistent with our hypothesis that tonic signaling and basal T-cell activation could potentiate T cells for rapid effector function. At the same time, sustained aerobic glycolytic activity is also associated with the acquisition of senescent phenotypes and a lack of potential for long-term persistence and memory formation (Kishton et al., 2017), consistent with the inability of rituximab-based CAR-T cells to achieve sustained control of tumor xenografts. This stands in stark contrast to the hybrid CAR-T cells that exhibit relatively low glycolytic flux but robust and sustained anti-tumor activity. A major challenge faced by CAR-T cells with intrinsically high metabolic rates is metabolic competition with tumor cells, which could further constrain the CAR-T cells’ ability to sustain their function (Chang et al., 2015). The Warburg effect, characterized by high rates of glucose uptake and lactate secretion despite the presence of oxygen, is a hallmark of tumor cells (Liberti and Locasale, 2016). In the tumor microenvironment, CAR-T cells and tumor cells must compete for the same limited supply of nutrients and contribute to the same potentially toxic acidification of the tumor microenvironment via lactate secretion (Fischer et al., 2007). This competition could have a disproportionately strong impact on the anti-tumor efficacy of rituximab-based CAR-T cells compared to other CAR-T cell variants with lower metabolic burdens.

Given that high metabolic rates are necessary to support robust effector T-cell function while memory phenotypes are conducive to long-term tumor control, an intriguing question is whether combining different CAR-T cells that target the same antigen—e.g., by co-administering rituximab CAR-T cells with RFR-LCDR.AA CAR-T cells either simultaneously or sequentially— would achieve greater therapeutic efficacy than administering either cell population alone. The use of multiple cell products could incur significant costs and technical complications compared to single-product administration. Nevertheless, such combinations may prove useful in conditions that have thus far resisted response to CAR-T cell therapy.

The structural modularity of CAR molecules supports the development of CAR-T cell therapies for a wide range of disorders, but the ability to rationally design CARs that yield predictably robust function *in vivo* remains elusive. The protein-engineering strategies and CAR-T cell characterization methods explored in this study offer a systematic approach to tune CAR signaling and quantitatively evaluate CAR-T cell phenotype and metabolism, supporting the engineering of next-generation CAR-T cells for refractory diseases that currently lack effective options.

## MATERIALS AND METHODS

### Construction of anti-CD20 scFvs and CARs

Plasmids encoding scFv sequences of rituximab and GA101 were generous gifts from Dr. Anna M. Wu (Zettlitz et al., 2017) (UCLA and City of Hope). DNA sequence encoding the ofatumumab scFv was codon optimized and synthesized by Integrated DNA Technologies (IDT; Coralville, IA). Plasmid encoding scFv derived from the leu16 monoclonal antibody (mAb) was a generous gift from Dr. Michael C. Jensen (Seattle Children’s Research Institute) (Jensen et al., 1998). V_L_ and V_H_ sequences of anti-GD2 scFv were identified from the 14g2a mAb (PDB code 4TUJ) using abYsis (Swindells et al., 2017). Anti-CD20 CARs were constructed by assembling an scFv (in V_L_-V_H_ orientation), an extracellular IgG4 hinge-CH2-CH3 spacer containing the L235E N297Q mutation (Hudecek et al., 2015), CD28 transmembrane and cytoplasmic domain, CD3ζ cytoplasmic domain, and a T2A “self-cleaving” sequence followed by a truncated epidermal growth factor receptor (EGFRt) with the MSCV backbone. EGFRt was used as a transduction and sorting marker. The abovementioned anti-CD20 CAR constructs were used as templates to generate CAR-HaloTag fusion proteins for microscopy imaging of CAR clustering.

### Cell line generation and maintenance

HEK 293T and Raji cells were obtained from ATCC. K562 cells were a gift from Dr. Michael C. Jensen (Seattle Children’s Research Institute). CD19^+^CD20^+^ K562 cells were generated as previously described (Zah et al., 2016). Luciferase-expressing CHLA-255 cell line (CHLA-255-Luc) was a gift from Dr. Shahab Asgharzadeh (Children’s Hospital of Los Angeles). CHLA-255-Luc-EGFP cells were generated by retroviral transduction of CHLA-255-Luc to express EGFP, and EGFP+ cells were enriched by fluorescence-activated cell sorting (FACS) on FACSAria (II) (BD Bioscience) at the UCLA Flow Cytometry Core Facility. HEK 293T cells were cultured in DMEM (HyClone) supplemented with 10% heat-inactivated FBS (HI-FBS; ThermoFisher). CHLA-255-Luc-EGFP cells were cultured in IMDM (ThermoFisher) with 10% HI-FBS. Primary human T cells, Raji, and K562 cells were cultured in RPMI-1640 (Lonza) with 10% HI-FBS. For CAR-T cells used in metabolomics studies, T cells were cultured in RPMI-1640 containing 2 g/L of 1,2-^13^C-glucose with 10% heat-inactivated dialyzed FBS (HI-dFBS).

### Retrovirus production and generation of human primary CAR-T cells

Retroviral supernatants were produced by transient co-transfection of HEK 293T cells with plasmids encoding anti-CD20 CAR or control constructs, and pRD114/pHIT60 virus-packaging plasmids (gifts from Dr. Steven Feldman), using linear polyethylenimine (PEI, 25 kDa; Polysciences). Supernatants were collected 48 and 72 hours later and pooled after removal of cell debris by a 0.45 μm membrane filter. Healthy donor blood was obtained from the UCLA Blood and Platelet Center. CD8^+^ T cells were isolated using RosetteSep Human CD8^+^ T Cell Enrichment Cocktail (StemCell Technologies) following manufacturer’s protocol. Peripheral blood mononuclear cells (PBMCs) were isolated from a Ficoll-Paque PLUS (GE Healthcare) density gradient. CD14^−^/CD25^−^/CD62L^+^ naïve/memory T cells (T_N/M_) were enriched from PBMCs using magnetic-activated cell sorting (MACS; Miltenyi). CD8+ T cells were used in early *in vitro* studies and in the *in vivo* study shown in Fig. 1B,C. T_N/M_ cells, which were more recently shown to exhibit clinical potential as a highly functional starting population for therapeutic T-cell manufacturing (NCT04007029; Zah et al., 2020), were used in later *in vivo* studies, including those shown in Figs. 2 and 4. Both CD8+ T cells and T_N/M_ cells were stimulated with CD3/CD28 Dynabeads (ThermoFisher) at a 3:1 cell-to-bead ratio on Day 0 (day of isolation) and transduced with retroviral supernatant on Day 2 and Day 3. Dynabeads were removed on Day 7. T cells were cultured in RPMI-1640 supplemented with 10% HI-FBS and fed with recombinant human IL-2 (ThermoFisher) and IL-15 (Miltenyi) every 2 days to final concentrations of 50 U/mL and 1 ng/mL, respectively. For CAR-T cells used in RNA-seq, ATAC-seq and metabolomic study, T cells were enriched for CAR^+^ expression by magnetic cell sorting via staining of EGFRt with biotinylated cetuximab (Eli Lilly; biotinylated in-house) followed by anti-biotin microbeads (Miltenyi). For RNA-seq and ATAC-seq, dead cells were depleted with a dead cell removal kit (Miltenyi) prior to enrichment of EGFRt^+^ population.

### Cytokine production quantification

Fifty thousand CAR+ T cells on Day 13 or 14 were incubated with 25,000 EGFP-expressing parental K562 (CD19^−^CD20^−^) or CD19^+^CD20^+^ K562 target cells at a 2:1 effector-to-target (E:T) ratio in 96-well U-bottom plate. To control for cell density while accounting for differences in transduction efficiency, untransduced T cells were added as necessary to reach the same number of total T cells per well. After a 48-hour co-incubation, cells were spun down at 300 xg for 2 min. Supernatant was harvested and cytokine levels were quantified by ELISA (BioLegend).

### Proliferation assay

T cells were stained with 1.25 μM CellTrace Violet (ThermoFisher) and 40,000 CAR^+^ T cells were seeded in each well in 96-well U-bottom plates with parental K562 or CD19^+^CD20^+^ K562 cells at a 2:1 E:T ratio. Untransduced T cells were added to wells as needed to normalize for differing transduction efficiencies and ensure the total number of T cells per well was consistent throughout. Cultures were passaged as needed, and CTV dilution was analyzed on a MACSQuant VYB flow cytometer after a 4-day co-incubation.

### Cytotoxicity assay with repeated antigen challenge

CAR^+^ T cells were seeded at 4 × 10^5^ cells/well in 24-well plate and coincubated with target cells at a 2:1 E:T ratio. Untransduced T cells were added to wells as needed to normalize for differing transduction efficiencies and ensure the total number of T cells per well was consistent throughout. Cell counts were quantified by a MACSQuant VYB flow cytometer every 2 days prior to addition of fresh target cells (2 × 10^5^ cells/well).

### Antibody staining for flow-cytometry analysis

EGFRt expression was measured with biotinylated cetuximab (Eli Lilly; biotinylated in-house), followed by PE-conjugated streptavidin (Jackson ImmunoResearch #016-110-084). CAR expression was quantified by surface epitope staining using Flag tag (DYKDDDDK tag, APC, clone L5, BioLegend #637308), HA (FITC, clone GG8-1F3.3.1, Miltenyi #130-120-722), or with anti-Fc (Alexa Fluor 488, Jackson ImmunoResearch #709-546-098). Antigen-independent activation-marker expression of CAR-T cells was evaluated by antibody staining for CD137 (PE/Cy7, clone 4B4-1, BioLegend #309818) and CTLA-4 (PE/Cy7, clone BNI3, BioLegend #369614) on Days 18 (i.e., 18 days after Dynabead addition and 11 days after Dynabead removal). Antibodies for CD45RA (VioGreen, clone T6D11, Miltenyi # 130-113-361), CD62L (APC, clone DREG-56, ThermoFisher #17-0629-42), PD-1 (APC, clone EH12.2H7, BioLegend #329908), and LAG-3 (APC, clone 3DS223H, ThermoFisher #17-2239-42) were used to evaluate T-cell subtype and exhaustion status. T-cell and tumor-cell persistence *in vivo* were monitored by antibody staining of retro-orbital blood samples. Samples were treated with red blood cell lysis solution (10X, Miltenyi) following manufacturer’s protocol. The remaining cellular content was stained with anti-human CD45 (PacBlue or PECy7, clone HI30, BioLegend #304029 or #304016) and biotinylated cetuximab, followed by PE-conjugated streptavidin. All samples were analyzed on a MACSQuant VYB flow cytometer (Miltenyi), and the resulting data were analyzed using the FlowJo software (TreeStar).

### Confocal microscopy

Jurkat cells transduced with CAR-HaloTag fusion protein were seeded at 10,000 cells per well in 50 μL RPMI-1640 + 10% HI-FBS in one well of a 48-well flat-bottom glass plate (MatTek). Scanning confocal imaging was acquired with a Zeiss LSM 880 laser scanning confocal microscope with AiryScan and a 63X 1.4 NA oil objective.

### *In vivo* studies

Six- to 8-week old NOD/SCID/IL-2Rγ^null^ (NSG) mice were obtained from UCLA Department of Radiation and Oncology. The protocol was approved by UCLA Institutional Animal Care and Used Committee. For evaluation of CD19 and CD20 CAR-T cells, each mouse was administered 0.5 × 10^5^ EGFP^+^ firefly luciferase (ffLuc)-expressing Raji cells by tail-vein injection. Six to nine days later, 1.5 × 10^6^ – 5 × 10^6^ CAR^+^ T cells or cells expressing EGFRt only (negative control) were injected via tail vein to tumor-bearing mice after confirming tumor engraftment. A second dose at 0.5 × 10^6^ and a third dose at 2 × 10^6^ of tumor cells were administered in a subset of studies as noted in the text and figures. In the study of GD2 CAR-T cells, each mouse was administered with 3.5 × 10^6^ CHLA-255-Luc-EGFP cells by tail-vein injection. Upon confirmation of CHLA-255 tumor engraftment (17 days post tumor injection), 2 × 10^6^ CAR^+^ T cells were injected via tail vein to tumor-bearing mice. Details of tumor dose, T-cell dose, tumor re-challenge were indicated in the text and figures. Tumor progression/regression was monitored with an IVIS Illumina III LT Imaging System (PerkinElmer). Blood samples were harvested via retro-orbital bleeding 3 days post T-cell injection and every 10 days thereafter. Mice were euthanized at the humane endpoint. Bone marrow, spleen and liver were collected after euthanasia. Tissues were ground and passed through a 100-μm filter followed by red-blood-cell lysis prior to flow-cytometry analysis. For ATAC-seq and RNA-seq experiments, NSG mice were engrafted with 0.5 × 10^6^ Raji cells 6 days prior to treatment with 2.85 × 10^6^ CAR-T cells. Nine days after T-cell injection, CAR-T cells were recovered from animal tissues (liver, spleen, cardiac blood, and bone marrow) and enriched for huCD45^+^ and EGFRt^+^ subpopulation by magnetic-activating cell sorting (MACS) prior to ATAC-seq/RNA-seq library construction.

### ATAC-seq library construction and data analysis

ATAC-seq libraries were constructed as previously described (Buenrostro et al., 2013; Corces et al., 2017). In brief, 30,000 – 50,000 viable T cells per mouse from MACS sort were washed once with PBS and lysed in 50 μL Resuspension Buffer (RSB) buffer (10 mM Tris-HCL, pH 7.4, 10 mM NaCl, 3 mM MgCl_2_) with 0.1% IGEPAL CA-630, 0.1% Tween-20, and 0.01% digitonin. Samples were washed with 1 mL RSB buffer containing 0.1% Tween-20 and centrifuged at 500 xg for 10 minutes at 4 °C. Pelleted nuclei were resuspended in 25 μL Tn5 transposition mix (12.5 μL 2X Tagment DNA buffer, 1.25 μL Tn5 transposase, and 11.25 μL sterile water; Illumina) and stored in a shaking incubator at 37°C and 500 RPM for one hour. Transposition reaction was purified with DNA Clean & Concentrator kit (Zymo Research). DNA fragments were PCR-amplified using NEB Q5 MasterMix and custom primers as previously described (Buenrostro et al., 2013). Libraries were size selected by AmPure beads (Beckman Coulter) and quantified by TapeStation. Libraries were sequenced on the Illumina NovaSeq S1 platform at the High Throughput Sequencing core at UCLA Broad Stem Cell Research Center with 50-bp paired-end reads. Fastq files from ATAC-seq were quality examined by FastQC (Linux, v0.11.8). Reads were processed by cutadapt (Linux, v1.18) to remove reads with low quality (quality score < 33) and to trim adapters. Trimmed reads were aligned to mm10 reference genome using Bowtie2 (Linux, v2.2.9) to eliminate contaminating reads from mouse cells. Non-murine reads were subsequently mapped to hg38 genome by Bowtie2, and sam files were converted to bam files by samtools (Linux, v1.9). Peaks were called independently for each replicate using MACS2 (Linux, v2.1.2) on the aligned reads, and subsequently merged by bedtools (v2.26.0). Reads assigned to each peak were counted by f*eatureCounts* function in subread (Linux, v1.6.3). To visualize chromatin accessible sites, peaks called from MACS2 were visualized in IGV (v2.8.0). Fold enrichments (calculated by MACS2) of peaks within –1 kb to 1 kb of the transcription start site (TSS) indicate accessibility of promoter regions.

### Bulk RNA-seq and gene set enrichment analysis (GSEA)

Total RNA was extracted from 200,000 – 700,000 MACS-sorted CAR-T cells using Qiagen RNeasy Plus Mini kit. mRNAs were isolated using NEBNext Poly(A) mRNA Magnetic Isolation Module (New England BioLabs). RNA-seq libraries were generated using NEBNext Ultra II Directional RNA Library Prep Kit (New England BioLabs) following manufacturer’s protocol. Fastq files from RNA-seq were quality-examined by FastQC (Linux, v0.11.8). Reads were processed by cutadapt (Linux, v1.18) to remove reads with low quality (quality score < 33) and to trim adapters. Trimmed reads were aligned to mm10 reference genome using Tophat2 (Linux, v2.1.0) to remove the contaminated reads from mouse cells. Non-murine reads were mapped to hg38 genome by Tophat2. Reads assigned to each gene were counted by *featureCounts* function in subread package (Linux, v1.6.3) with ensembl 38 gene sets as references. Genes without at least 8 reads mapped in at least one sample were considered below reliable detection limit and eliminated. Read counts were normalized by Trimmed Mean of M-values method (TMM normalization method in edgeR running on R v3.6.3) to yield RPKM (reads per millions per kilobases) values, and differential expression was calculated using the package edgeR. Gene ontology analysis was performed using GSEA software (v4.1.0, Broad Institute) and BubbleGUM (v1.3.19) (Spinelli et al., 2015). Expression values of differentially expressed genes were input to the program and using a curated list of 2493 T-cell–relevant gene sets selected from current MSigDB gene sets. Heatmaps for differentially expressed genes were generated using *pheamap* and *ggplot2* packages in R (version 3.6.3). Volcano plots were generated using *ggplot2* in R (version 3.6.3).

### Metabolite extraction and analysis

Cells were initially cultured in RPMI-1640 supplemented with 10% HI-FBS. At 24 to 72 hours before metabolite extraction, culture media were changed to RPMI containing 10% FBS or dialyzed FBS (dFBS) with 2 g/L 1,2-^13^C-glucose. Cell culture media were collected from each cell line every 24 hours to evaluate nutrient uptake and consumption. Four volumes of 100% HPLC-grade methanol were added to one volume of media and centrifuged at 17,000 xg and 4°C for 5 minutes to precipitate cell debris. Clear supernatants were harvested and analyzed by liquid chromatography followed by mass spectrometry (LC-MS). To provide accurate estimation of nutrient uptake and consumption, partial media change was performed every 24 hours to avoid nutrient depletion. Twenty-four or 72 hours after switching to labeled-glucose media, intracellular metabolite extraction was performed as previously described (Bennett et al., 2008; Park et al., 2019). In brief, cells were transferred onto nylon membrane filters (0.45 μm; Millipore) and vacuumed to remove media. Each filter was quickly soaked in 400 μL cold extraction solvent (HPLC-grade acetonitrile:methanol:water 40:40:20, v/v) in one well of a 6-well plate. The plates were incubated at –20°C for 20 minutes. Cell extracts were subsequently transferred to 1.7 mL microcentrifuge tubes and centrifuged at 17,000 xg in 4°C for 5 minutes. Supernatants were lyophilized and reconstituted in HPLC-grade water normalized to total cell count (50 μL per 1 million cells). Both methanol-treated media samples and intracellular metabolite extracts were analyzed by reversed-phase ion-pairing liquid chromatography (Vanquish UPLC; Thermo Fisher Scientific) coupled to a high-resolution orbitrap mass spectrometer (Q-Exactive plus Orbitrap; Thermo Fisher Scientific) at the Molecular Instrumentation Center (MIC) in UCLA. Metabolites were identified by comparing mass-to-charge (m/z) ratio and retention time to previously validated standards. Samples were detected in both negative-ion mode and positive-ion mode. Negative-ion mode was separated into two subgroups—nlo and nhi—to obtain data with m/z ratio from 60 to 200 and 200 to 2000, respectively. LC-MS data were processed using Metabolomic Analysis and Visualization Engine (MAVEN) (Clasquin et al., 2012). Labeling fractions were corrected for the naturally occurring abundance of ^13^C. Concentration of metabolites in culture media was quantified at 24 and 72 hr by normalizing ion counts from LC-MS measurement to controls with known concentrations. Uptake and secretion rates were calculated by subtracting sample concentration from fresh media and normalizing to viable cell count (positive values indicated secretion and negative values indicated uptake). A mole balance was performed to account for media change from cell cultures. Calculation accounted for 10–20% of media evaporation every 24 hours.

### Statistical Analysis

Statistical tests including two-tailed, unpaired, two-sample Student’s *t* test and log-rank Mentel-Cox test were performed using GraphPad Prism V8. One-way ANOVA test for differential gene analysis in RNA-seq was performed with *glmQLFTest* function in edgeR.

## ACKNOWLEDGMENTS

This work was supported by the Alliance for Cancer Gene Therapy and the Cancer Research Institute (grants to Y.Y.C.), as well as the UCLA Council on Research Faculty Research Grant (grant to J.O.P.). A.L. was supported by the UCLA Biotechnology Training in Biomedical Sciences and Engineering Program (NIH/NIGMS T32-GM067555-15). We thank Drs. Anna M. Wu and Kirstin A. Zettlitz for helpful discussions on anti-CD20 scFv sequences. We thank Dr. Willy Hugo and Katherine M. Sheu for helpful guidance and discussions on RNA-seq data analysis. We thank Wallace Wennerberg, Jaimie Chen, Sabah Rahman, Brenda Ji, Paul Ayoub, Duo Xu, and Zheng Cao for their technical assistance. We thank Suhua Feng, Marco Morselli, Shawn Cokus and Mahnaz Akhavan (UCLA BSCRC BioSequencing Core Facility), and UCLA Molecular Instrument Center for their technical support. This work was performed with instrumentation and computational resources from the Eli and Edythe Broad Center of Regenerative Medicine and Stem Cell Research – Molecular, Cell, and Developmental Biology Microscopy Core, the UCLA Jonsson Comprehensive Cancer Center (JCCC) Flow Cytometry Core Facility supported by the National Institutes of Health (P30 CA016042), and the Hoffman2 Shared Cluster provided by the UCLA Institute for Digital Research and Education’s Research Technology Group.

## AUTHOR CONTRIBUTIONS

Y.Y.C. and X.C. designed the research and wrote the manuscript; J.O.P. designed and supervised metabolomics experiments; X.C., M.K., A.L., E.S., A.S., A.A., Y.D., D.N., J.O.P. performed experiments; X.C., M.K., A.L., X.M., J.O.P. and Y.Y.C. performed data analysis; L.C. contributed technical discussion on alanine insertion.

