## Supplementary Figures for "Rational Tuning of CAR Tonic Signaling Yields Superior T-Cell Therapy for Cancer"

### SUPPLEMENTAL FIGURES

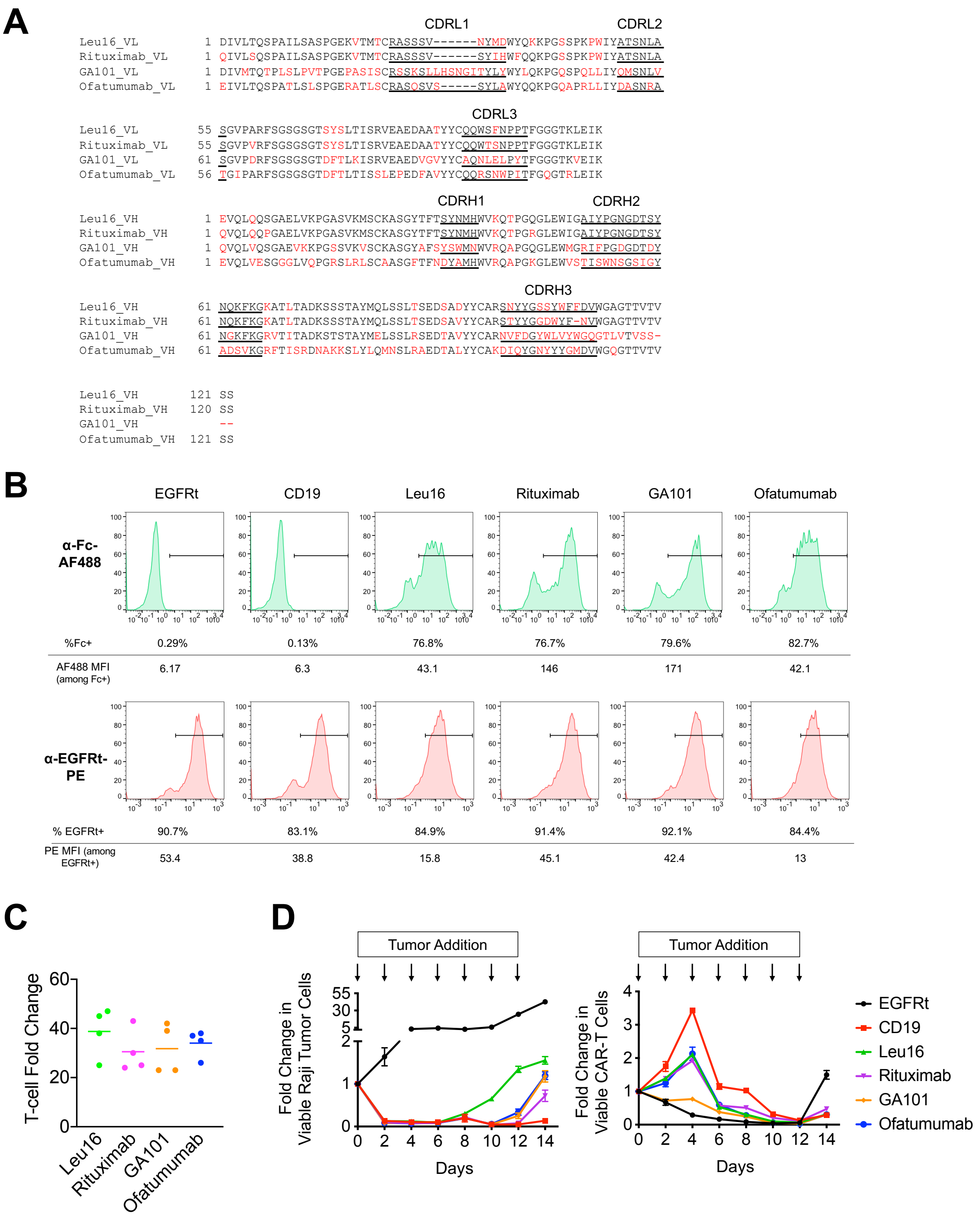


**Figure S1. Panel if CD20 CARs exhibit similar characteristics *in vitro***

(A) Alignment of leu16, rituximab, GA101 and ofatumumab scFv sequences using T-Coffee (Notredame et al., 2000).

(B) CAR-T cell transduction efficiency as quantified by CAR surface expression (top) and transduction-marker expression (bottom), which were detected via antibody staining of the CAR's IgG4 extracellular spacer (Fc) and EGFRt, respectively. Median florescence intensity

(MFI) and % positive of each antigen staining were noted below flow cytometry histograms. Results are representative of three independent experiments from three different healthy donors.

(C) CD20 CAR-T cell proliferation during *ex vivo* culture with exogenous cytokine IL-2 and IL15. Fold change in total viable cell count between Day 2 and Day 14 are shown. Each data point represents one donor; data for 4 donors per construct are shown. No statistical difference between any of the donors was detected by unpaired, two-tailed, two-sample Student's *t* test.

(D) CAR-T cell cytotoxicity and proliferation upon repeated antigen challenge. CD20 CAR-T cells were challenged with Raji tumor cells at a 2:1 effector-to-target (E:T) ratio every two days, and the number of viable Raji and CAR-T cell was quantified by flow cytometry. Data shown are the means of technical triplicates with error bars indicating ± 1 standard deviation (S.D.). Results are representative of three independent experiments from three different healthy donors.

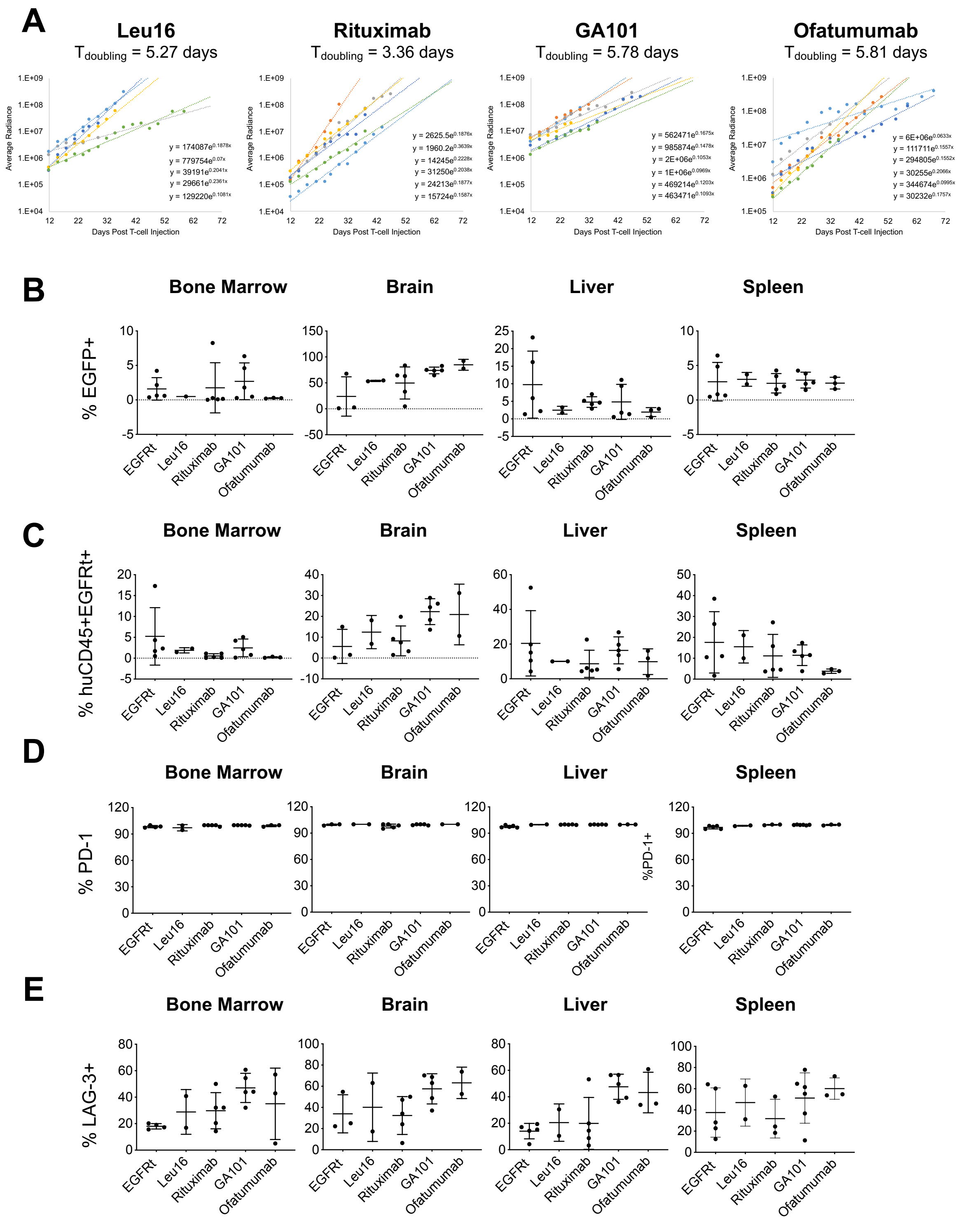


**Figure S2. Tumor progression and post-mortem CD20 CAR-T cell characterization in Raji xenograft model**

(A) NSG mice were engrafted with firefly-luciferase–expressing Raji cells and treated with CD20 CAR-T cells as described in Figure 1B. Tumor burden was measured by bioluminescence imaging, and the tumor growth rate between Day 12 post T-cell injection and the humane end point was quantified by fitting exponential regression curves to each individual animal's tumor progression data. The average doubling time ($T_{doubling}$) for tumor signal among the mice (n = 6) in each group is shown above each plot.

(B, C) Frequency of tumor cells (B) and human $CD45^{+}EGFRt^{+}$ cell (B) in bone marrow, brain, liver, and spleen collected from mice at the time of euthanasia, as quantified by flow cytometry.

(D,E) Frequency of PD-$1^{+}$ (D) and LAG-$3^{+}$ (E) cells among human $CD45^{+}EGFRt^{+}$ cell in bone marrow, brain, liver, and spleen collected from mice at the time of euthanasia, as quantified by flow cytometry. No statistical difference between any of the donors was detected by unpaired, two-tailed, two-sample Student's *t* test for all data shown in panels (B) through (E).

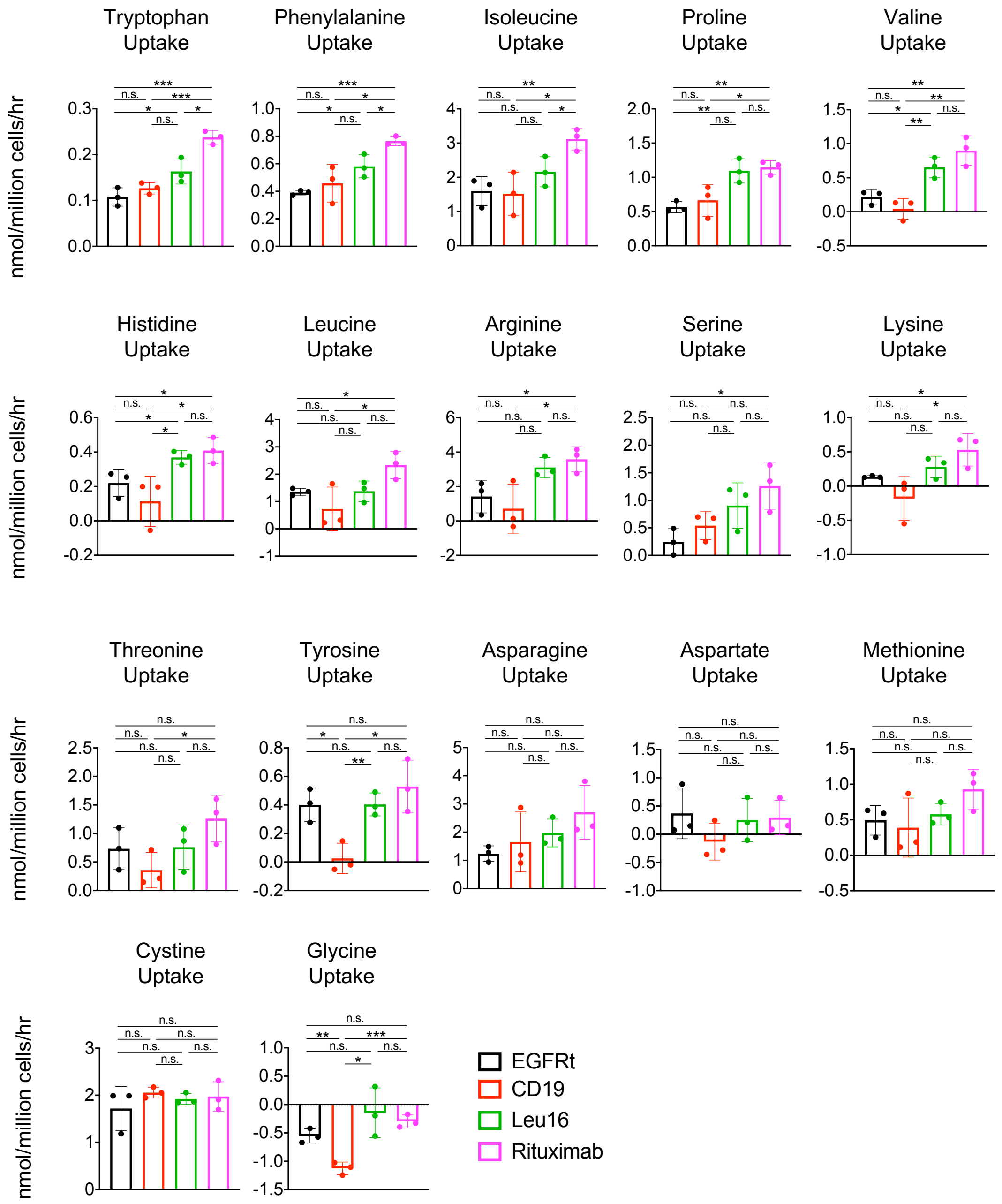


**Figure S3. Rituximab CAR-T cells are more metabolically activate than other CD20 CAR-T cells**

Uptake of amino acids and other nutrients by CAR-T cells cultured for 24 hours in RPMI supplemented with 10% heat-inactivated dialyzed fetal bovine serum (HI-dFBS), IL-2, and IL-15. Data bars indicate the means of technical triplicates ± 1 S.D. Results are representative of three independent experiments from three different healthy donors. **p*<0.05, ***p*<0.01, ****p*<0.001, n.s. not statistically significant. Results in this figure are from the same experiment as in Figure 1F.

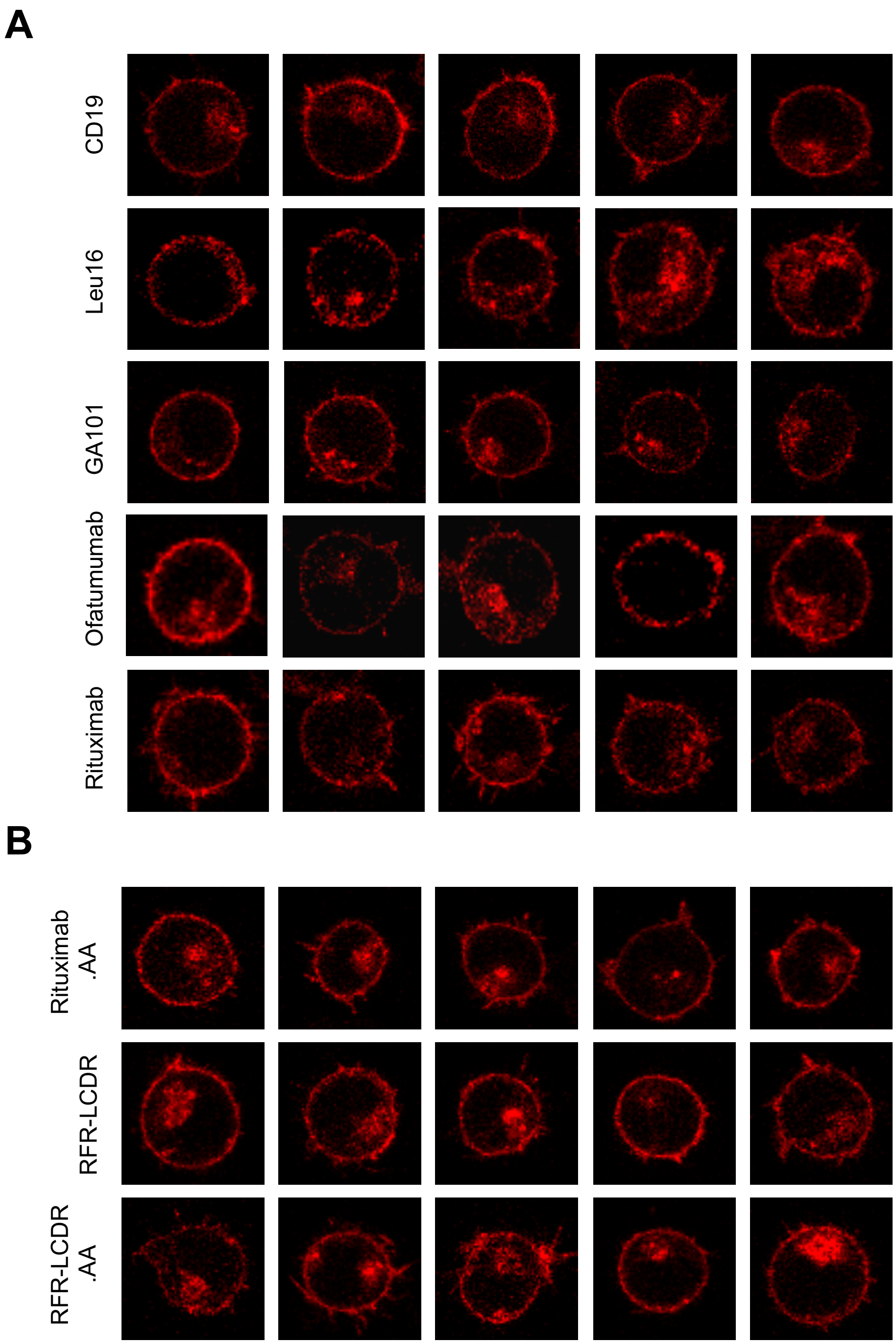


**Figure S4. CD20 CARs are uniformly distributed on T-cell surface in the absence of antigen engagement**

Jurkat cells transduced with CAR-HaloTag fusion proteins were stained with the red fluorescent dye tetramethylrhodamine (TMR) and imaged by confocal microscopy. CAR molecules are uniformly distributed on cell surface in the absence of antigen stimulation. **(A)** Original CD20 CAR panel. **(B)** CD20 CARs incorporating alanine insertion and/or scFv sequence hybridization.

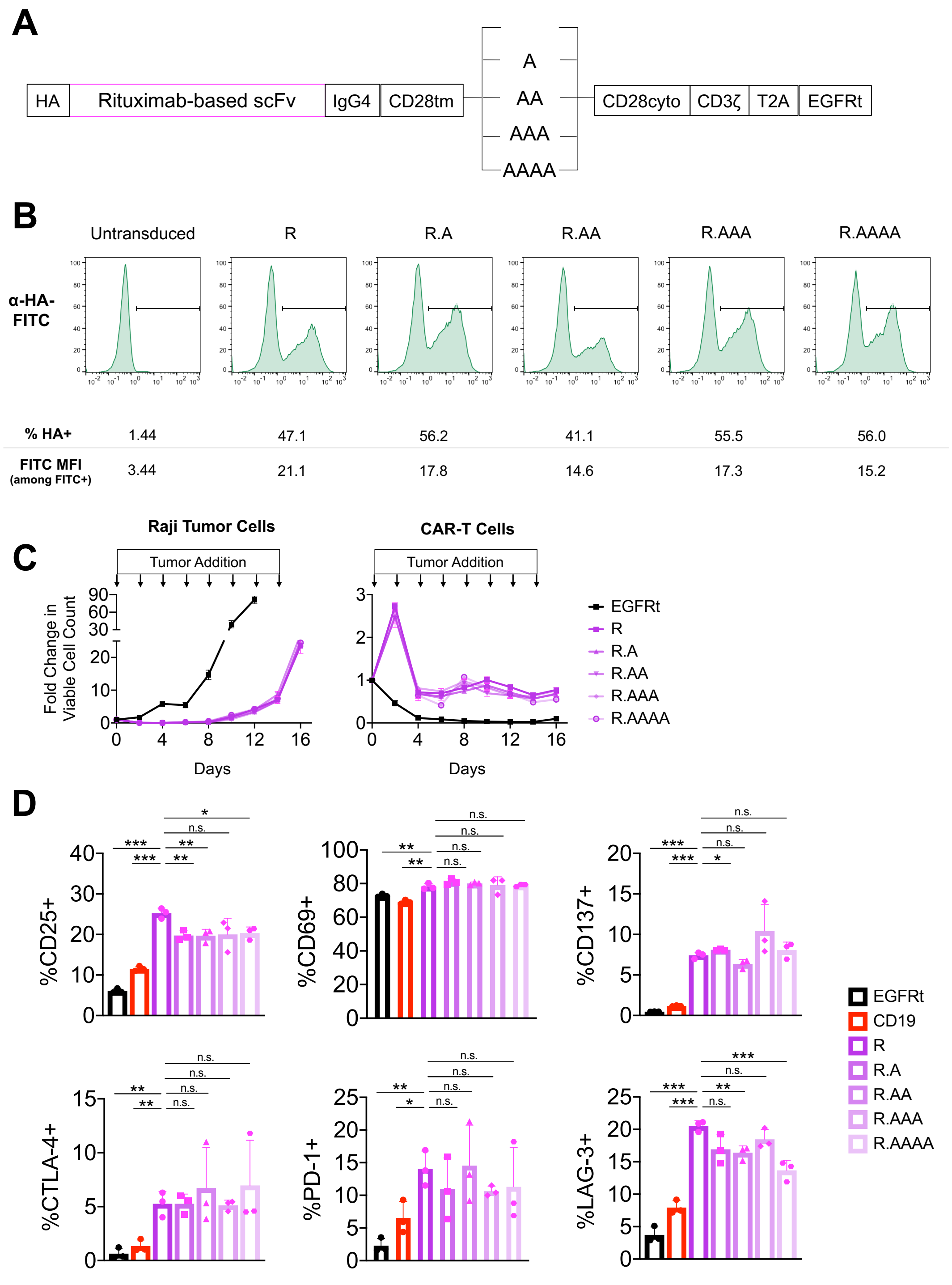


**Figure S5. Rituximab CAR-T cells with torsional reorientation exhibit similar *in vitro* cytotoxicity and antigen-independent activation-marker expression**

(A) Schematic of rituximab-based CAR constructs with zero to four alanines inserted between the CD28 transmembrane and cytoplasmic domains.

(B) CAR surface expression was quantified by antibody staining of HA tag fused to the N-terminus of each CAR. Data are representative of three independent experiments from three different healthy donors.

(C) CAR-T cell cytotoxicity and proliferation upon repeated antigen challenge. CD20 CAR-T cells were challenged with Raji tumor cells at a 2:1 E:T ratio every two days, and the number of viable Raji and CAR-T cell was quantified by flow cytometry. Data shown are the means of technical triplicates with error bars indicating ± 1 standard deviation (S.D.). Results are representative of three independent experiments from three different healthy donors.
(D) Activation- and exhaustion-maker expression were evaluated 11 days post Dynabead removal, without CD20 antigen stimulation. Data bars indicate the means of technical triplicates ± 1 S.D. Results are representative of three independent experiments from three different healthy donors. Unless otherwise noted, p values were determined by unpaired two-tailed Student's *t* test; *$p$<0.05, **$p$<0.01, ***$p$<0.001, n.s. not statistically significant.

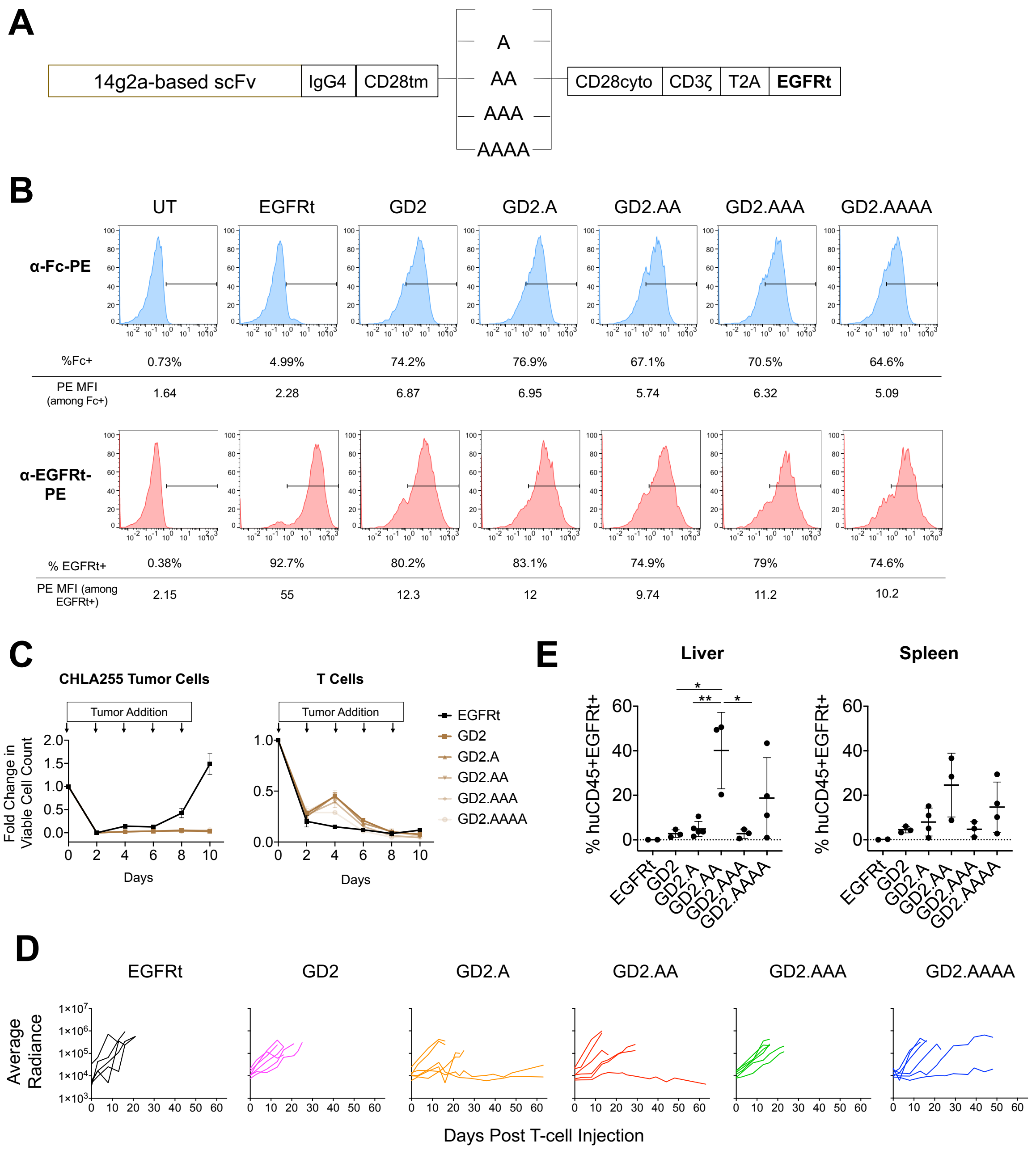


**Figure S6. GD2 CAR-T cells with torsional reorientation exhibit differential tumor control *in vivo* despite same *in vitro* performance**

(A) Schematic of GD2 CAR constructs with zero to four alanines inserted between the CD28 transmembrane and cytoplasmic domains.

(B) CAR surface expression (top) and transduction efficiency (bottom) were quantified by antibody staining of Fc and EGFRt.

(C) CAR-T cell cytotoxicity and proliferation upon repeated antigen challenge. GD2 CAR-T cells were challenged with CHLA-255 tumor cells at a 2:1 E:T ratio every two days, and the number of viable Raji and CAR-T cell was quantified by flow cytometry. Data shown are the means of technical triplicates with error bars indicating ± 1 standard deviation (S.D.).

(D,E) NSG mice were injected intravenously with 3.5 x $10^6$ firefly-luciferase expressing CHLA-255 cells followed by 2 x $10^6$ GD2 CAR-T cells 17 days later.
(D) Tumor progression was monitored by bioluminescence imaging.
(E) Frequency of human $CD45^+EGFRt^+$ cell in liver and spleen collected from mice at the time of euthanasia was quantified by flow cytometry. *$p$<0.05, **$p$<0.01, ***$p$<0.001, n.s. not statistically significant.

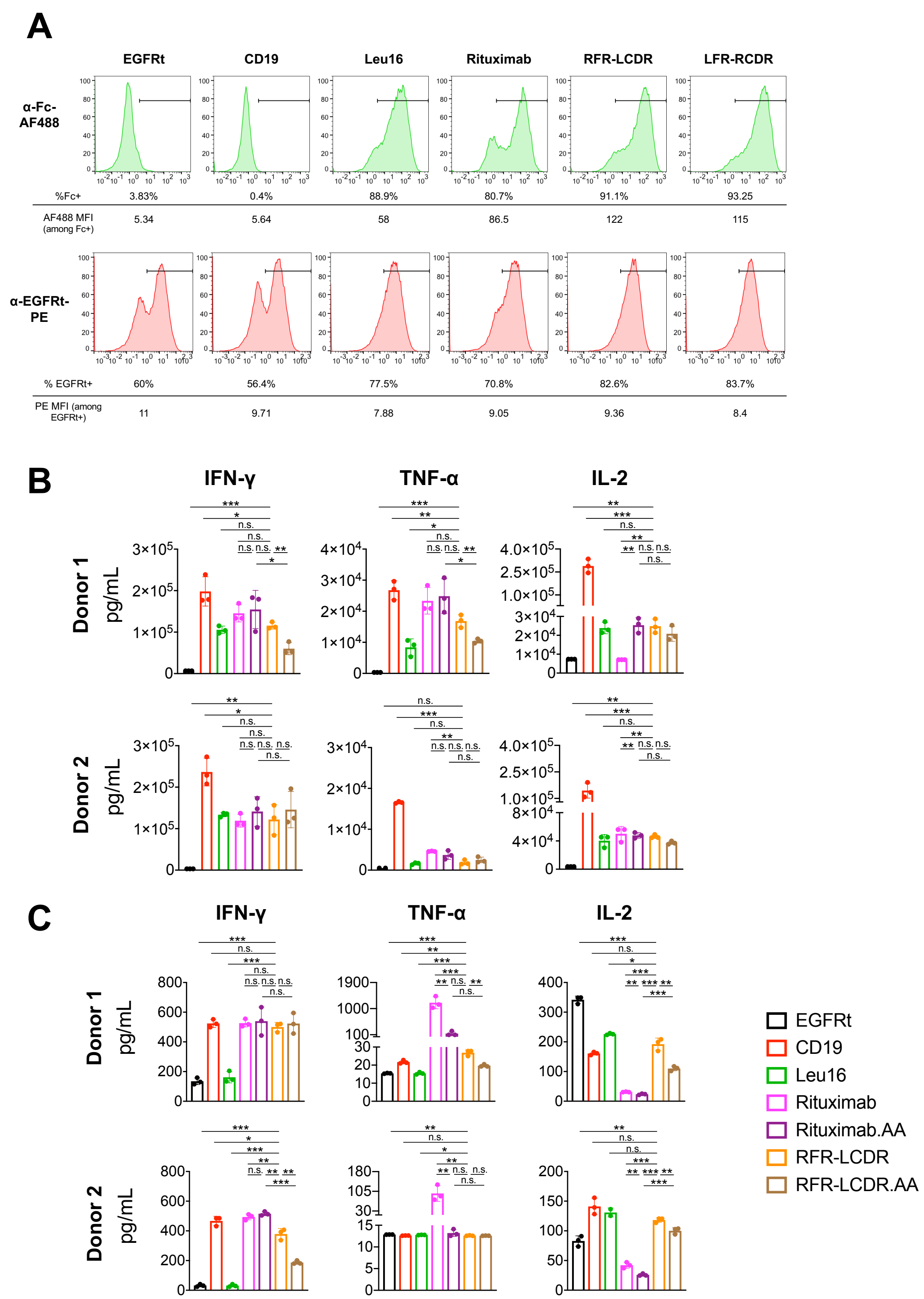


**Figure S7. *In vitro* characterization of hybrid CAR-T cells**

(A) CAR surface expression (top) and transduction efficiency (bottom) were quantified by antibody staining of Fc and EGFRt. Results are representative of three independent experiments from three different healthy donors.

(B,C) IFN-γ, TNF-α, IL-2 production in the presence (C) and absence (D) of $CD19^+/CD20^+$ K562 target cells. **p*<0.05, ***p*<0.01, ****p*<0.001, n.s. not statistically significant.

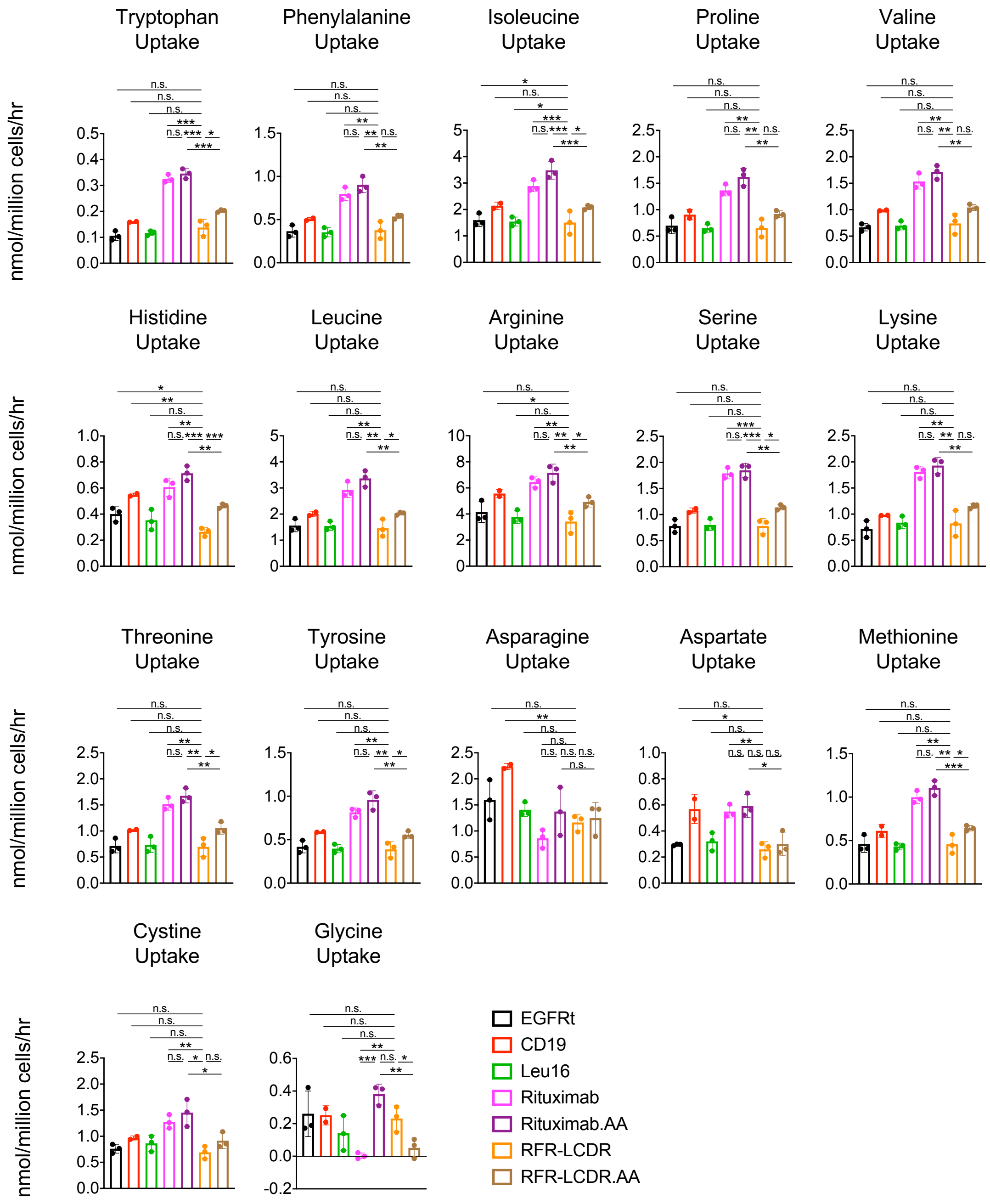


**Figure S8. Hybrid CAR-T cells exhibit reduced metabolic activities compared to rituximab CAR-T cells**

Uptake of amino acids and other nutrients by CAR-T cells cultured for 72 hours in RPMI supplemented with 10% heat-inactivated dialyzed fetal bovine serum (HI-dFBS), IL-2, and IL-15. Data bars indicate the means of technical triplicates ± 1 S.D. Data are representative of three independent experiments from three different healthy donors. *$p$<0.05, **$p$<0.01, ***$p$<0.001, n.s. not statistically significant. Results in this figure are from the same experiment as in Figure 3E.

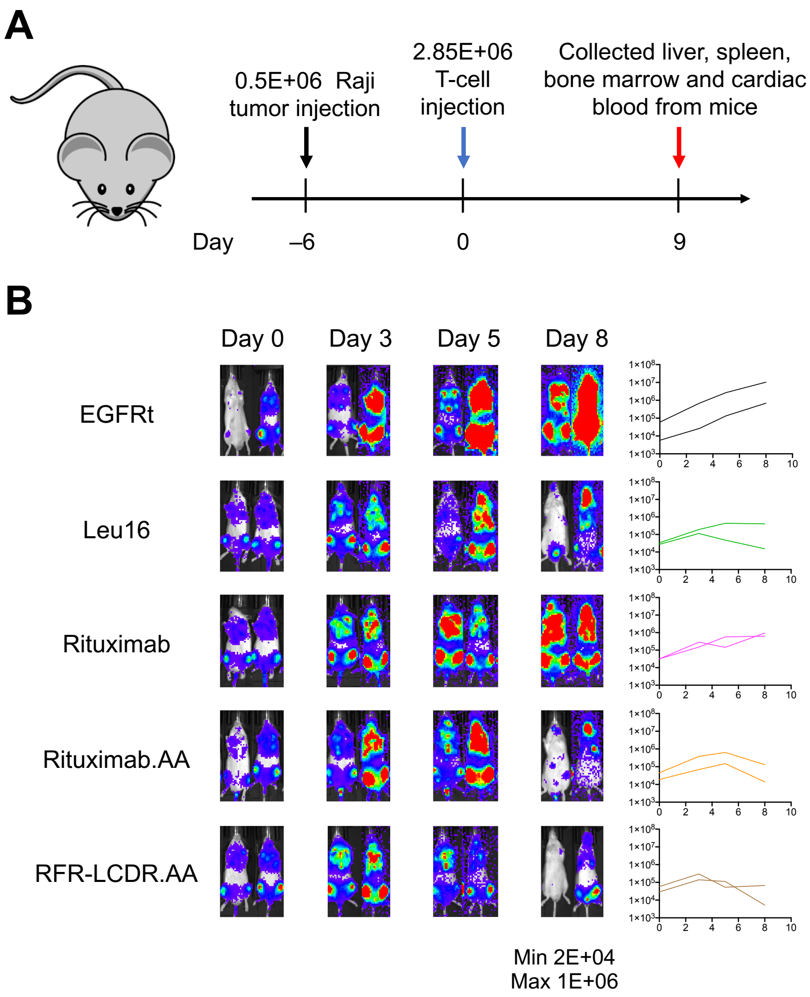


**Figure S9. CAR-T cell harvest from tumor-bearing mice for transcriptomic and epigenetic profiling**

NSG mice were injected intravenously with 0.5 x $10^6$ firefly-luciferase expressing Raji cells followed by 2.85 x $10^6$ of $CAR^+$ T cells 6 days later. $CAR^+$ T cells were collected from tumor-bearing mice 9 days after T-cell injection (n = 2 mice per group). Only a small number of mock-transduced (EGFRt-only) T cells were recovered from mice, consistent with lack of T-cell expansion in the absence of antigen recognition. However, this poor cell recovery led to low read counts in RNA-seq and ATAC-seq that precluded reliable data analysis. As a result, these samples were excluded from the analyses shown in Figures 5 and 6 and Supplementary Figure 10.

(A) Schematic of *in vivo* experiment.

(B) Tumor progression as monitored by bioluminescence imaging.

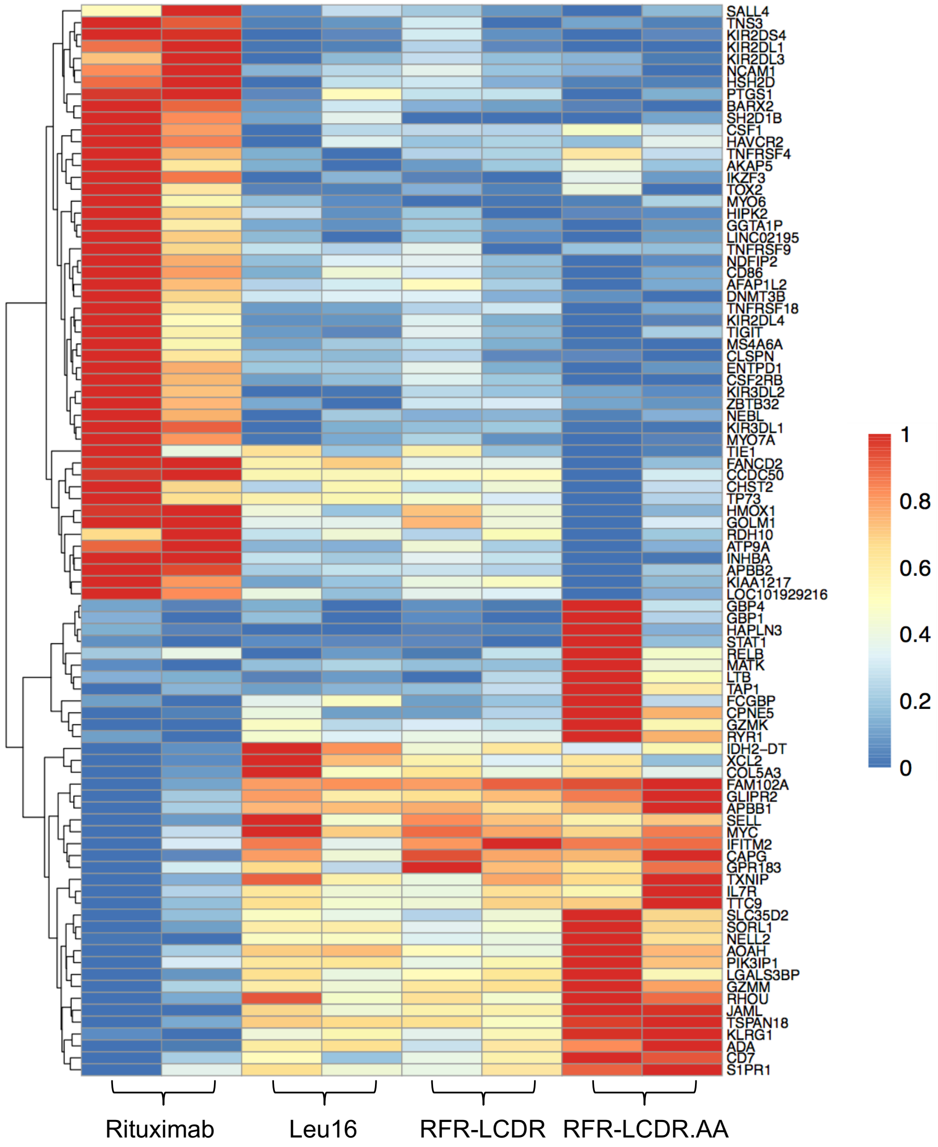


**Figure S10. RNA-seq reveals drastic transcriptomic differences across CD20 CAR-T cell variants recovered from tumor-bearing mice**

Heatmap of differentially expressed genes (FDR < 0.05) in ANOVA comparison of Leu16, Rituximab, RFR-LCDR, RFR-LCDR.AA CAR-T cells. Each column represents one mouse, and two biological replicates were analyzed for each treatment group. Each row is scaled to a maximum of 1 and minimum of 0 to highlight relative expression of each gene.

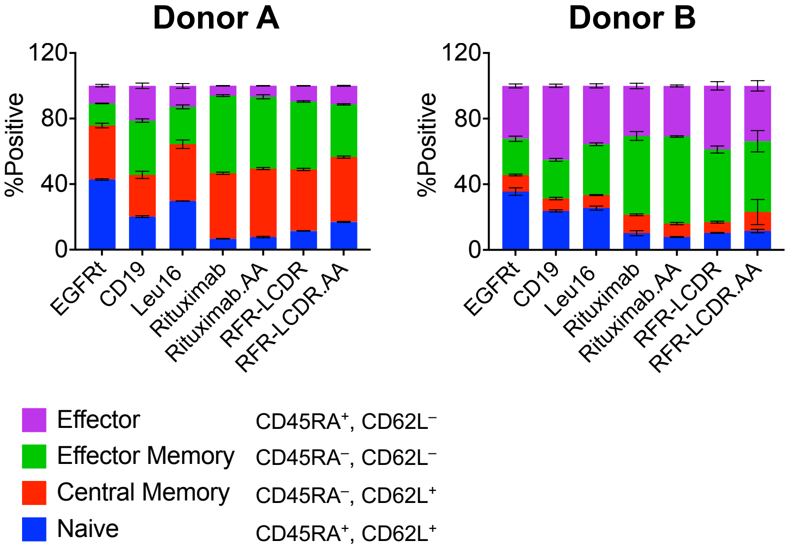


**Figure S11. T-cell subset distribution does not vary significantly among CAR-T cells in *ex vivo* culture**

The subtype of CAR-T cells in *ex vivo* culture was determined by CD45RA and CD62L staining in the absence of antigen stimulation. $CD45RA^{+}CD62L^{+}$, $CD45RA^{-}CD62L^{+}$, $CD45RA^{-}CD62L^{-}$, and $CD45RA^{+}CD62L^{-}$ indicate naïve, central memory, effector memory and effector cell types, respectively.
